# A delay in sampling information from temporally autocorrelated visual stimuli

**DOI:** 10.1101/656850

**Authors:** Chloe Callahan-Flintoft, Alex O. Holcombe, Brad Wyble

## Abstract

Much of our world changes smoothly in time, yet the allocation of attention is typically studied with sudden changes – transients. When stimuli change gradually there is a sizeable lag between when a cue is presented and when an object is sampled (Carlson, Hogendoorn, & Verstraten, 2006; Sheth, Nijhawan & Shimojo, 2000). Yet this lag is not seen with rapid serial visual presentation (RSVP) stimuli where temporally uncorrelated stimuli are presented (Vul, Kanwisher & Nieuwenstein 2008; Goodbourn & Holcombe, 2015). These findings collectively suggest that temporal autocorrelation of a feature paradoxically increases the latency at which information is sampled. This hypothesis was tested by comparing stimuli changing smoothly in time (autocorrelated) to stimuli that change randomly. Participants attempted to report the color coincident with a visual cue. The result was a smaller selection lag for the randomly varying condition relative to the condition with a smooth color trajectory. Our third experiment finds that the increase in selection latency is due to the smoothness of the color change after the cue rather than extrapolated predictions based on the color changes presented before the cue. Together, these results support a theory of attentional drag, whereby attention remains engaged at a location longer when features are changing smoothly. A computational model provides insights into neural mechanisms that might underlie the effect.

## Introduction

The visual system allows for the selection and prioritization of certain pieces of information in our visual field over others. As our visual input is constantly changing, due to scene changes as well as head and eye movements, the visual system needs the ability to make this selection not only in space but also in time before the information is gone.

Classic studies of the time-course of attention involved a target stimulus presented at a variable time or SOA (stimulus onset asynchrony) after the sudden onset of a cue at the target location. The target was immediately masked after it was presented, and the logic was that the minimum SOA needed for accurate performance provides an estimate of the time for attention to arrive at and sample the cued location (Nakayama & Mackeben, 1989; Muller & Rabbit, 1989). These studies found that performance rapidly increased as SOA increases to reach a peak around 80-120 ms after the cue. This work has been influential and guided the current understanding of the time needed to shift attention and sample a stimulus. However, it is not clear how straightforwardly such results apply to a continuous stream of potential target stimuli, as is often encountered in real-world scenes.

Extraction of a target at a particular time from a continuous stream of input is a nontrivial problem because the representation of a given stimulus is distributed across cortical areas with distinct processing latencies. Thus, there may be uncertainty regarding the relative time of a cue and the potential target stimuli. Moreover, the visual system may tend to group together stimuli over time, creating a potential segmentation problem, particularly when the features of those stimuli are temporally autocorrelated. For the remainder of this paper the term *smooth* will be used as a term for temporal autocorrelation of stimulus features.

One line of research using smoothly changing features displayed a set of clock faces whose hand smoothly changed throughout the trial. Participants were asked to report the hand position of the clock at the time of a cue. On average, participants reported the clock hand position presented about 130 ms after the cue (Carlson, Hogendoorn, & Verstraten, 2006). Using a similar clock paradigm, this magnitude of lag has also been seen in tasks exploring divided attention (Hogendoorn, Carlson, VanRullen, & Verstraten, 2010) and attentional shifts (Chakravarthi & VanRullen, 2011). It is not limited to the sampling of position. Displaying a disk that changed smoothly in color, Sheth, Nijhawan & Shimojo (2000) flashed another colored disk on the opposite side of fixation. Participants on average reported the color of the changing disk as it was approximately 400 ms after the flashed cue disk. Delays were also found for objects changing smoothly in luminance (37 ms), spatial frequency (83 ms) and entropy (95 ms).

Substantial lags for sampling from a stream of stimuli are not always found. Vul, Kanwisher & Nieuwenstein (2008) and Goodbourn & Holcombe (2015) used one or two simultaneous streams of letters and flashed a cuing circle around one of the streams. Participants attempted to report the letter that was presented in that stream at the time of the cue. The average delay estimated was very small, 25 ms or less in both cases. A notable difference with the studies that found longer lags is that those with longer lags used stimuli that changed smoothly over time, while Vul et al. (2008) and Goodbourn & Holcombe (2015) presented a sequence of unrelated letters that changed abruptly from one to the next. Thus, it may be the case that sharp visual transients influence the latency of attention sampling. Previous studies have found that presenting a transient visual signal at the location of a changing stimulus improves temporal sampling whereas endogenous shifts of attention were not as effective (Holcombe & Cavanagh, 2008).

Based on this difference in findings between these paradigms we propose a theory of *attentional drag*, which posits that a temporal autocorrelation in a visual feature extends the duration of attentional engagement elicited by a cue in order to extract useful information from an object. Thus, attention gets dragged along in time by a temporally autocorrelated feature dimension, which increases the latency to disengage, and results in a delayed feature selection. Conversely, when features change abruptly, as in the case of a conventional RSVP stream of letters, the salient featural change after the cue leads to an earlier disengagement of attention and thereby an earlier selection. Thus, the attentional drag theory proposes that transients can decrease the latency to sample information in part by making it easier for attention to disengage.

By comparing two conditions, one where colors change randomly at a regular interval and another where they change smoothly, Experiment 1 shows empirically that selection can happen immediately in response to a cue in the random condition, but is delayed by about 100ms in the smooth condition. Critically, in both conditions the presentation rate of the stimuli is the same, the only difference is the temporal autocorrelation of colors.

In Experiment 2 these findings are extended to show that increasing the similarity between successive colors further extends the attentional window. Finally, Experiment 3 tested whether these results could be the product of feature extrapolation based on the trajectory prior to the flash and find that this explanation is unable to account for the results.

Together, the experimental and modeling work presented here demonstrate an influence of featural change in time on information extraction. The neurocomputational model provided in the discussion demonstrates how a simple attentional system can exhibit prolonged selection latencies. In a nutshell, the prolonged latencies reflects a feedback loop between attention-related neurons and sensory neurons. The sensory neurons have overlapping tuning curves that result in greater activation in the case of a smoothly changing stimulus, which in combination with the recurrence with attention neurons causes longer-lasting activation.

## Experiment 1

### Methods

For all experiments presented, the stimuli, code for running the paradigms, data, and the code for running all of the analysis and generating the figures of this paper have been uploaded to the Open Science Framework (OSF) (https://osf.io/hujwb/).

#### Participants

Twenty-five undergraduates from the Pennsylvania State University subject pool filled out informed consent forms before participating in this study. Participants were between 18 and 23 years old and had normal or corrected to normal vision.

#### Stimuli & Apparatus

Stimuli were presented using a 16-inch CRT monitor (1024×768, 75 Hz refresh rate) placed 63.5 cm away from participants’ headrest, using Psychtoolbox functions (Brainard, 1997). The stimuli for this experiment consisted of two colored dots, presented on either side of a fixation cross (3° of visual angle separation between the center of each dot and the fixation). Each dot subtended 2.7° of visual angle. Four color rings (sets of colors that each form a closed path through color space) were generated for this experiment. The values of each ring were calculated by first setting the a, b coordinates (in L*a*b color space) for each ring’s center ([0, 0], [50, 0], [20, 50], [-40, 30]). The radius of each ring was set to 60 and the L parameter for each ring was set to 70. Given these parameters, 360 points were calculated along each ring (Figure 1B). The monitor was not color-calibrated and thus the actual L*a*b values differ, but the property that the colors formed a closed smooth path was preserved. The RGB values of each color ring and the code used to generate them have been posted to the OSF page cited above. The starting color of each disk as well as the direction of change through the color ring was randomly selected at the start of each trial. Thus there were eight color trajectories to reduce the participants’ ability to predict the color that would appear, although this will be shown not to matter in a subsequent experiment.

**Figure 1.**
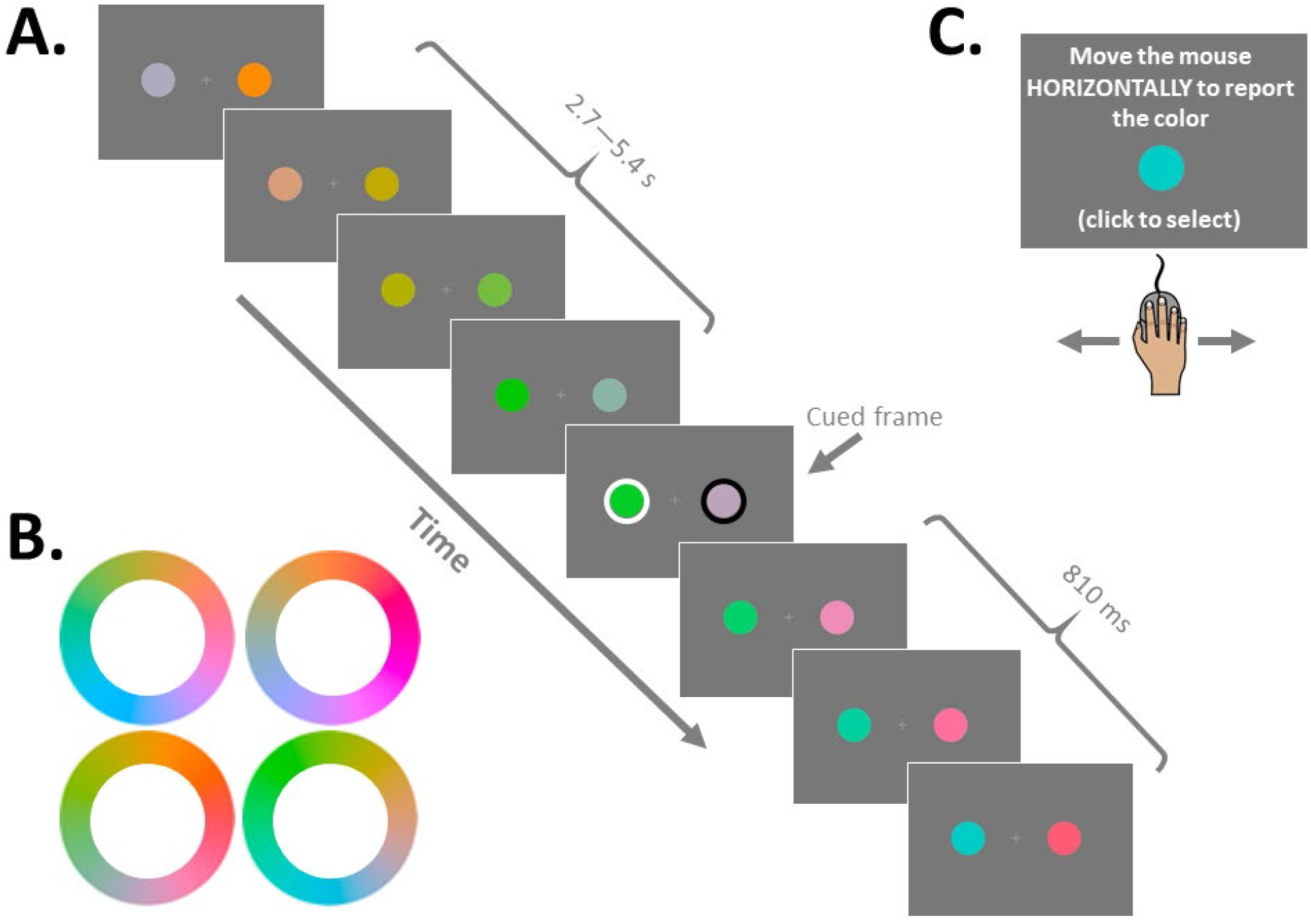
Experimental paradigm for Experiments 1, 1b, 2 & 2b. A) Participants fixated on a cross in the middle while two disks changed color (see Methods for details). A cue (white or black ring) flashed around one disk during the stream as a distractor disk flashed around the other for 27 ms. The disks then continued to change until the end of the trial. Here, sample frames of the Smooth condition are depicted. B) Four possible color rings. Two rings (one for each disk) was selected randomly at the start of each trial. C) At the end of the trial participants report the color of the disk at the time of the cue by moving the mouse horizontally to make the test circle smoothly change color. Participants click the mouse to select the color that best matches memory.

#### Procedure

Participants were asked to keep their eyes on a fixation cross in the middle of the screen while monitoring a changing color disk on both sides. Participants were told to remember the color of the disk when the cue (a white or black ring, counterbalanced across participants) flashed around it. Between 2,700 and 5,400 ms after the stimuli first appeared, the cue ring simultaneously onset around one of the disks and a distractor ring (the opposite color of the cue, white or black depending on the participant) appeared around the other disk for 27 ms. The timing of the cue, and the side on which the cue appeared, was randomized across trials. The two disks continued to change for another 810 ms after the offset of cue and distractor ring (Figure 1).

At the end of every trial a report screen instructed participants to move the mouse horizontally to change the color of a test disk to match that of the cued disk as it was at the time of the cue. The test disk was presented at the center of the screen. As the participants moved the mouse, the test disk changed colors through the cued disk’s color ring at a rate of 2 degrees per pixel. When the participant found the color that best matched their memory, he or she clicked the mouse to select it. Afterwards participants were given a score from 0 to 100 in which a score of 100 indicated that they had reported the exact color presented at the time of the cue and score of 0 meant that they had reported the color 180 degrees away from the target color. The score did not indicate the direction of error.

There were two conditions in this experiment each with 44 trials that were intermixed with in block. The first condition was the *Smooth* condition where the color of the disks advanced 16 degrees every 108 ms through the selected color ring. The second condition was the *Random* condition where the color disks changed pseudo-randomly every 108 ms. The selection of the next color was randomly chosen from the color ring of that disk with the constraint that the new color had to be at least 30 degrees away from the previous color. Thus, in both conditions color information changed at the same rate but the relationship in color space across time points differed.

### Results

The serial position error was calculated the same way for all experiments. On each trial the difference was taken between the reported color and the cued color as well as every color presented in the seven positions before and after the cue. The serial position error then was determined by the minimum difference (i.e. the color which was closest to the reported value) and translated into milliseconds using the SOA. The trial was excluded if the reported value was an equal distance from the two closest colors presented (i.e. a tie). Additionally, if this minimum difference between the reported color and the color of the closest matching stimulus was greater than 16 degrees of the 360 degrees of the color circle, that trial was excluded from analysis. The rationale was that such reports are especially likely to be guesses or gross misperceptions of the color. This second exclusion criteria was chosen a priori as a compromise between the Random condition, which requires no such criteria, and the Smooth condition which does in order to avoid edge effects, where the histograms bins at the edges of the minimum color response distribution (i.e. +/- 7 serial positions) would be assigned any guess responses from other portions of the color wheel. To check for robustness, additional analyses were performed varying the threshold for both Experiments 1 & 1b. The results and figures are included in the supplemental materials.

In order to test whether the difference in selection latency between two conditions was significantly different from zero, a permutation analysis was conducted. On every permutation, participants’ Smooth and Random condition position errors were randomly shuffled into two bins. The average position error of these two bins was then computed and the difference between those averages was recorded. This was repeated ten thousand times, building a null distribution of mean differences that would have resulted if there were no difference between the two conditions. The real mean difference between the Smooth and Random condition was compared to this permuted null distribution. The percentile location of the real mean difference in the permuted null distribution is the p-value^1^. In order to calculate a confidence interval around this mean difference, a stratified bootstrapping of the data was performed (to account for the uneven number of trials per participant due to the exclusion criteria). Ten thousand bootstrapped samples were drawn, taking the difference between means of each sample.

On average, 5 trials (*SD* = 3) were excluded from the Smooth condition and 7 trials (*SD* = 2) were excluded from the Random condition, per participant. The Smooth condition had a selection latency that was delayed by 101 ms relative to the Random condition (Figure 2). The difference in selection between the two conditions was significantly greater than zero, *p* <.001, 95% CI = [76, 135].

**Figure 2.**
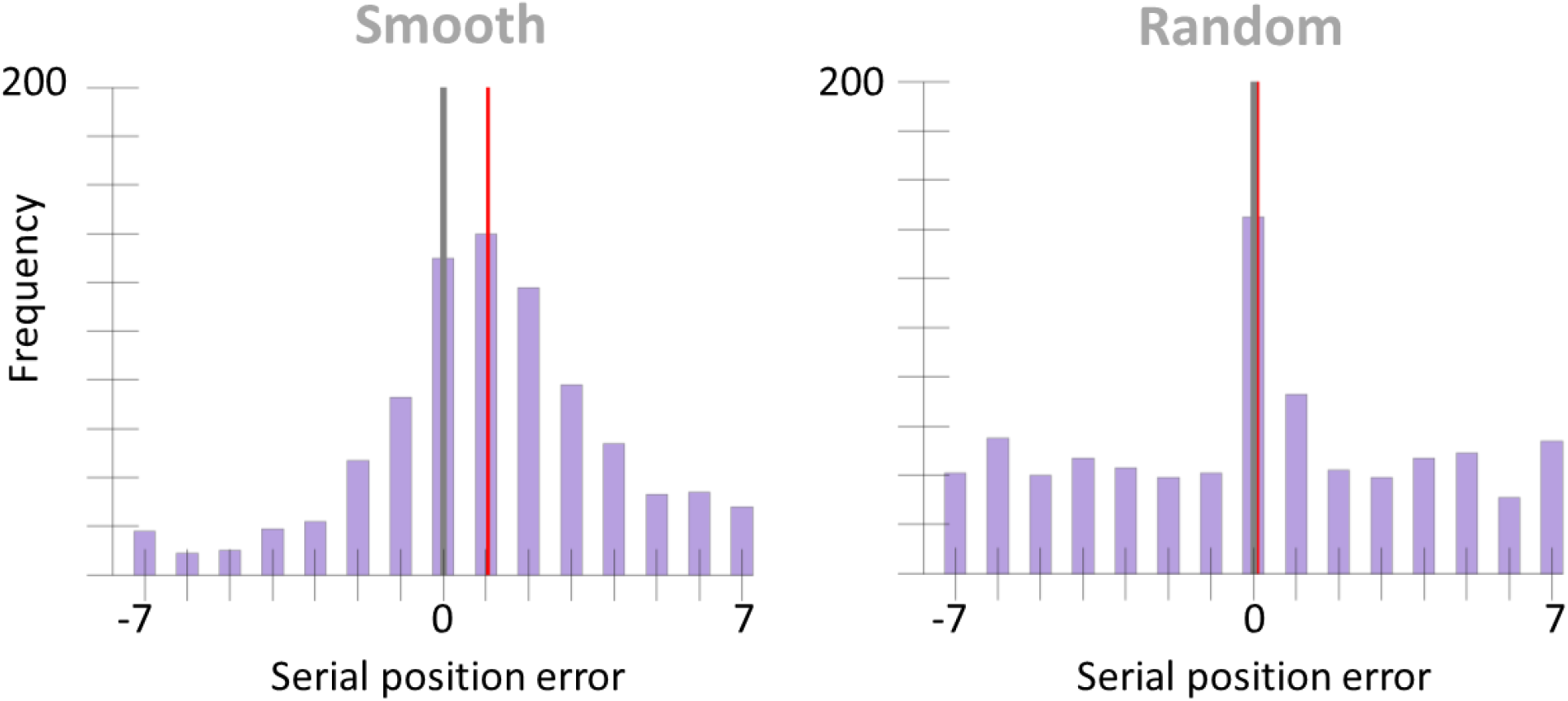
Serial position error histograms of Experiment 1’s Smooth and Random conditions. Error has been discretized by bucketing trials based on the presented color that was closest to the reported color (see Results section). The grey vertical line indicates the position of the cued color (position zero). The red vertical line marks the condition mean.

## Experiment 1b

Experiment 1b is a replication of Experiment 1, conducted to assess reproducibility of the results of Experiment 1.

### Methods

An independent sample of 25 participants were collected for this study. The exact same stimuli and procedure was used as in Experiment 1.

### Results

The same analysis technique was used here as in Experiment 1. Five trials (*SD* = 3) in the Smooth condition and 8 trials (*SD* = 3) in the Random condition were excluded from analysis on average per participant. The Smooth condition had a selection latency that was delayed by 113 ms relative to the Random condition (Figure 3). This difference was significant, *p* < .001, 95% CI = [78, 137].

**Figure 3.**
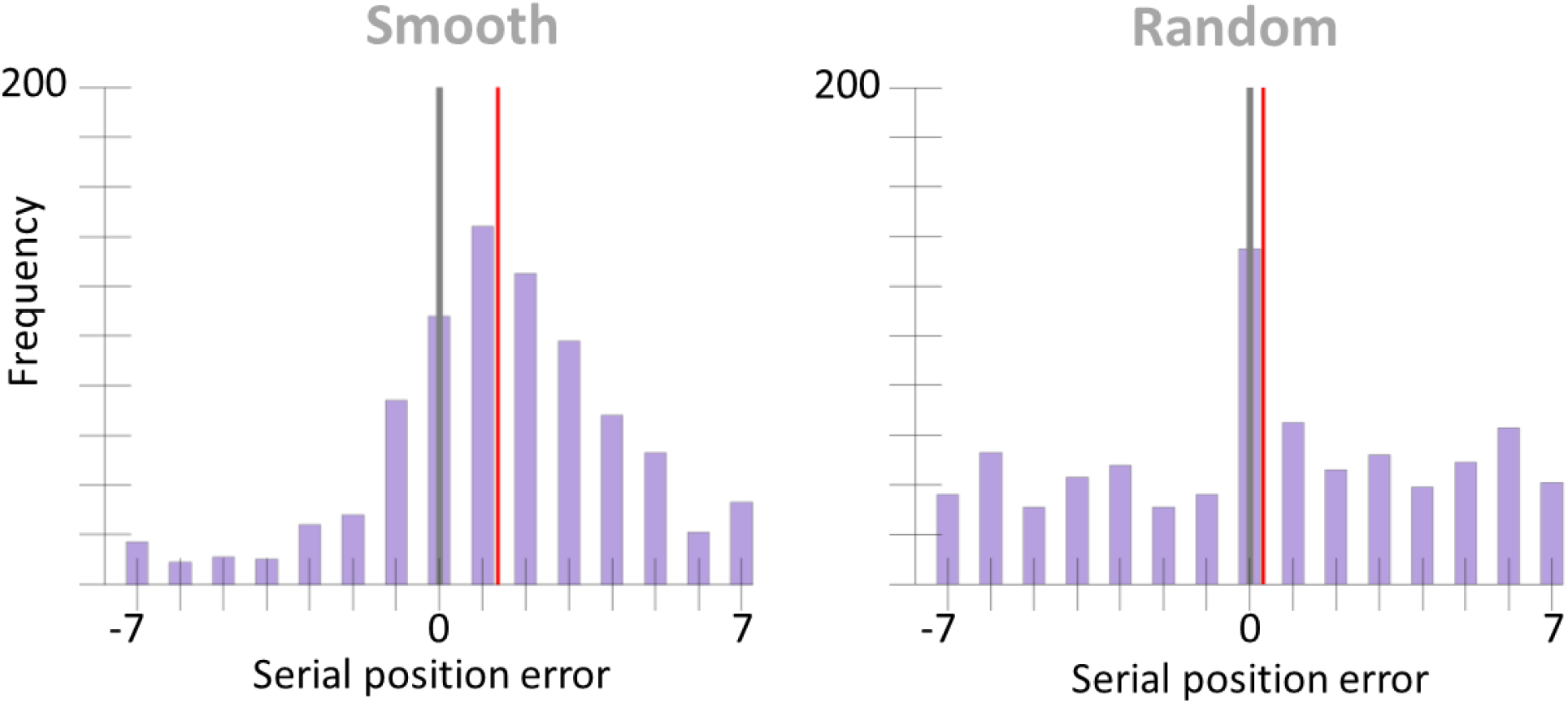
Serial position error histograms of Experiment 1b’s Smooth and Random conditions. The grey vertical line marks the cued color position and the red vertical line marks the condition mean.

### Discussion of Experiment 1 and 1b

The Random condition in both experiments yielded a more rapid attentional selection latency than the Smooth condition. These results suggest that something about the smooth changes leads to a later selected feature value than when stimuli are jumping randomly through feature space (i.e. the Random condition). This could be the product of extended attentional engagement induced by the featural autocorrelation. If attention is dragged along by temporal autocorrelation, then increasing the correlation between colors from one time point to the next should further increase the duration of attentional drag. To test this, Experiment 2 was run comparing the Smooth condition of Experiment 1 to a condition where the colors on successive frames are more similar to one another.

## Experiment 2

In Experiment 2 the Smooth condition of Experiment 1 was compared to a condition where the color trajectory was even smoother. The attentional drag theory predicts this smoother condition will result in a longer latency of color selection.

### Methods

#### Participants

Twenty-five undergraduates volunteered from the Pennsylvania State University subject pool to participate in this study. All participants were between 18 and 23 years old with normal or corrected to normal vision. Informed consent was obtained for each participant prior to the study in accordance with the IRB office of Penn State University.

#### Stimuli & Apparatus

The same stimuli used in Experiment 1 were used in Experiment 2.

#### Procedure

A procedure similar to Experiment 1’s was used in Experiment 2. Participants again monitored two color disks, one on either side of fixation, and were asked to report the color of the disk at the time a cue ring flashed around it. The same cueing and report method used in Experiment 1 were used here. Instead of Smooth and Random, the two conditions were Smooth_coarse and Smooth_fine. The Smooth_coarse condition here is the same as the Smooth condition of Experiment 1 where colors advanced 16 degrees every 108 ms along the color ring. In the Smooth_fine condition the color of the disks advanced along the color ring by 4 degrees every 27 ms. Importantly, in both conditions the average rate of change through color space was the same but finer time steps were taken in the Smooth_fine condition. As in Experiment 1, there were 44 trials per condition, intermixed within block.

### Results

As there were 4 times as many colors presented in the Smooth_fine condition as the Smooth_coarse, the number of positions (or color values presented) in the analysis before and after the cue was 28 in the Smooth_fine condition as opposed to 7 in the Smooth_Coarse. Similarly, since the Smooth_fine condition presents colors closer to one another along the color ring, the analysis for both conditions was restricted to only trials where the minimum distance between a reported color and presented colors was 8 degrees in this experiment. This criterion was chosen as a compromise in order to apply the same trial exclusion procedure to both conditions. A second analysis, included in the supplemental results, lowers the criterion to 4 degrees (ideal for the Smooth_fine condition). This naturally results in the exclusion of a greater number of trials, but it yields the same findings as those presented here (Figure S3). The same permutation analysis and bootstrapping method was used in this experiment as was described in the results of Experiment 1.

An average of 6 trials (*SD* = 3) from the Smooth_coarse condition and 12 trials (*SD* = 3) from the Smooth_fine condition were excluded from analysis by the exclusion criteria described above. The Smooth_fine condition had a significant delay in selection latency of 32 ms compared to the Smooth_coarse condition, *p* = .03, 95% CI [7, 58] (Figure 4).

**Figure 4.**
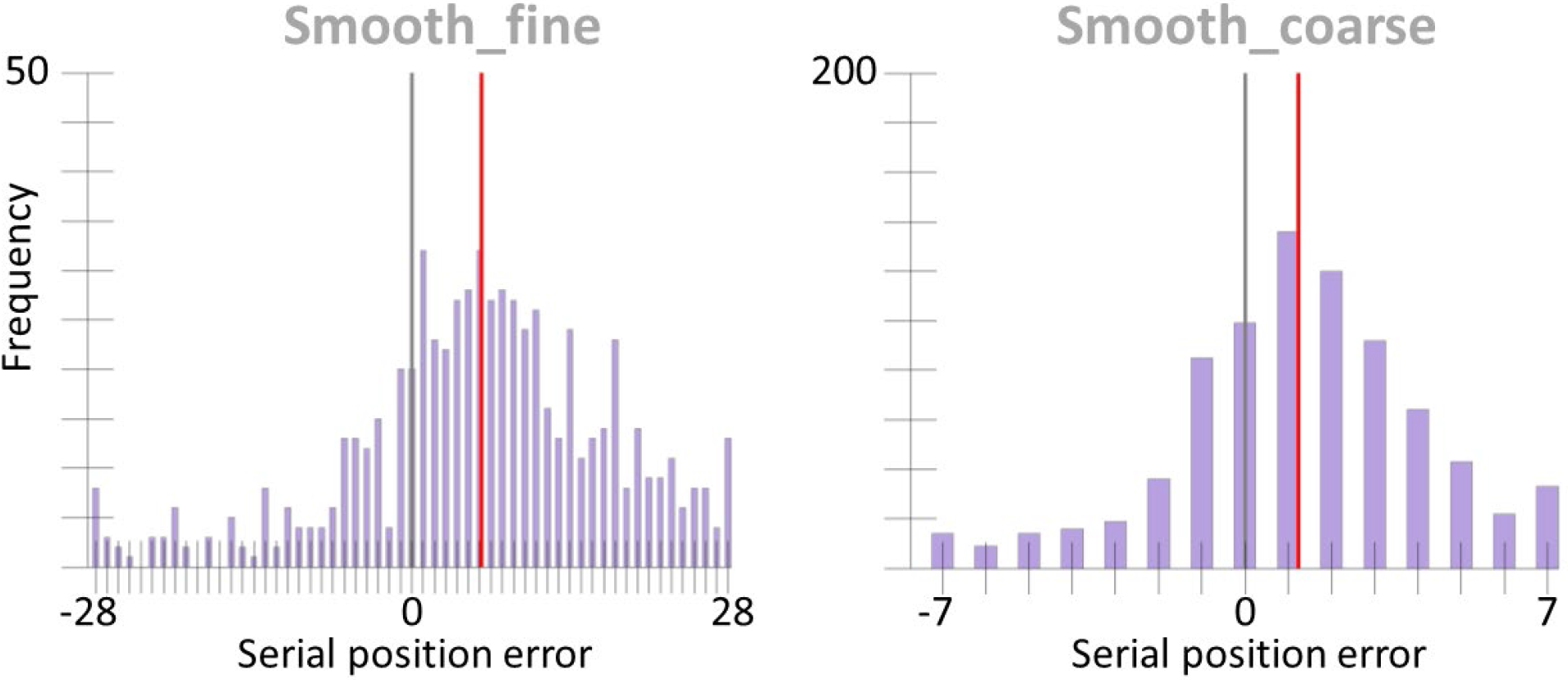
Serial position error histograms of Experiment 2’s Smooth_fine and Smooth_coarse conditions. Note there are four times the number of positions in the Smooth_fine condition as there are in the Smooth_coarse condition for the same window of time since colors updated in the Smooth_fine condition at four times the rate of that in the Smooth_coarse. The grey vertical line marks the position of the cued color. The red vertical line marks the condition mean.

## Experiment 2b

Experiment 2b was run as a replication of Experiment 2.

### Methods

An independent sample of 25 undergraduates from the Pennsylvania State University subject pool participated in this replication experiment. The exact same stimuli and procedure was used in Experiment 2b as was used in Experiment 2.

### Results

Per participant, an average of 6 trials (*SD* = 3) from the Smooth_coarse condition and 12 trials (*SD* = 3) from the Smooth_fine condition were excluded from analysis. The data were analyzed in the same way as were the data of Experiment 2. Experiment 2b replicated the delay in selection latency, 75 ms, between the Smooth_fine and Smooth_coarse condition, *p* < .001, 95% CI [8, 34] (Figure 5). As in Experiment 2, a second analysis was run with a more conservative exclusion criterion (only allowing a maximum of 4 degrees between a reported color and the color values presented), which found similar results, as shown in the supplemental results (Figure S4).

**Figure 5.**
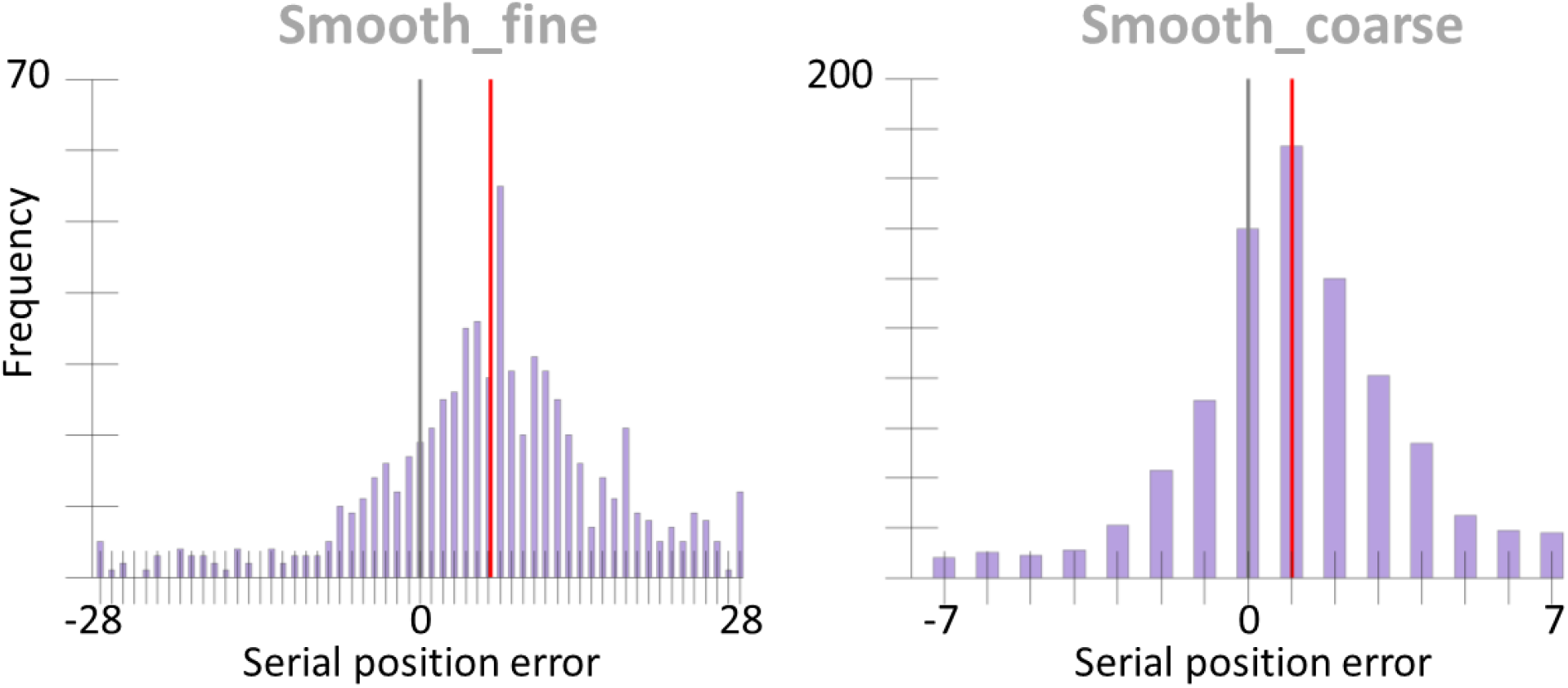
Serial position error histograms of Experiment 2b’s Smooth_fine and Smooth_coarse conditions. The grey vertical line marks the position of the cued color. The red vertical line marks the condition mean.

### Discussion of Experiment 2 and 2b

The results of Experiment 2 showed a significant reduction in the average selection latency from the Smooth_fine to Smooth_coarse presentation and this finding was supported by a replication. This result supports the hypothesis that smooth feature changes leads to prolonged attentional engagement and causes a later selection of feature information.

An alternative explanation for the combined results of Experiments 1 & 2 could be that instead of the increased disruption of smoothness from Smooth_fine to Smooth_coarse to Random presentation, it is actually the decrease in predictability across conditions that is producing the selection latency reduction. The increased jump in color space from one update to the next in the Smooth_coarse condition compared to the Smooth_fine condition may make it more difficult for the visual system to extrapolate the next color. If the visual system is sampling from a generated internal representation instead of the current stimulus input, this increased difficulty might reduce how far into the future the system can predict and result in what appears to be an earlier selection latency. This effect would similarly be seen again in comparing Smooth to Random selection in Experiment 1, as the Random condition does not offer a predictable change in color through time. However, it would be very counterintuitive for prediction effects to be the cause of these latency differences, since predictability is typically assumed to reduce errors rather than increase them.

To test the influence of predictability, Experiment 3 showed disks that changed in either a Smooth or Random pattern prior to the cue and switched to other pattern after the cue to test whether smoothness before or after the cue influenced selection latency.

## Experiment 3

Experiment 3 tested whether the selection latency differences seen in the previous two experiments were a result of the visual system generating predictions of the color trajectory. The theory would be that in generating these predictions, when the cue triggers attention, the color sampled is from the internal representation (a future value) which leads to an appearance of selecting information later in time. To test this a similar paradigm to the previous two was used but this time the colors changed smoothly either before or after the cue, and randomly otherwise. The internal model theory would predict that when the color changes are smooth before the cue, error distributions will be shifted forwards along the before-cue trajectory as color reports are based on predictions. Note, that in this scenario, the color trajectory that would have followed after the cue was never actually presented to the participant. Conversely, when participants are first presented with randomly changing colors and then a smooth change after the cue, this theory predicts less of a shift in the error distribution as the system is unable to generate predictions prior to the onset of attention.

### Method

#### Participants

A sample of 20 undergraduates age 18-23 with normal to correct to normal vision were used for this experiment with the same recruitment and consent procedure as outlined in Experiments 1 & 2.

#### Stimuli & Apparatus

The same stimuli used in Experiments 1 & 2 were used in this experiment.

#### Procedure

In this experiment participants again maintained fixation on a cross in the middle of the screen and monitored two changing colored disks. Participants were told to report the color of the disk at the time of the cue. The same cueing and reporting method used in the previous two experiments were used here.

In Experiment 3 there were two conditions. In the Smooth-to-Random condition the color disks changed color in accordance with the Smooth presentation style used in Experiment 1 (16 degrees along the color ring every 108 ms). At the time of the cue, the presentation style changed to random where colors were pseudo-randomly presented every 108 ms (with the same constraints as outlined in Experiment 1). In the Random-to-Smooth condition the color disks changed as in the Random condition of Experiment 1. At the time of the cue the disks began to change in the Smooth presentation style, beginning the color trajectory from the color presented at the time of the cue.

### Results

This experiment tested whether the latency reduction in Experiment 1 between the Smooth and Random condition was due to the visual system selecting a future prediction at the time of the cue based on the smooth change presented previously. If this is the case, participants should report colors further along a color trajectory when the color was presented smoothly prior to the onset of attention (the cue) even if the smoothness of that trajectory ended after the cue (as in the Smooth-to-Random condition). Conversely, participants should rely on the veridical colors presented when there is no smoothness to base prediction on, resulting in a selection around the time of the cue for the Random-to-Smooth condition. To test this we measured the distance between the reported value and smooth trajectory, as it would have appeared if the presentation style was constant throughout the trial (Figure 6). Again, the same permutation analysis and bootstrapping method was used in this experiment as was described in the results of Experiment 1.

**Figure 6.**
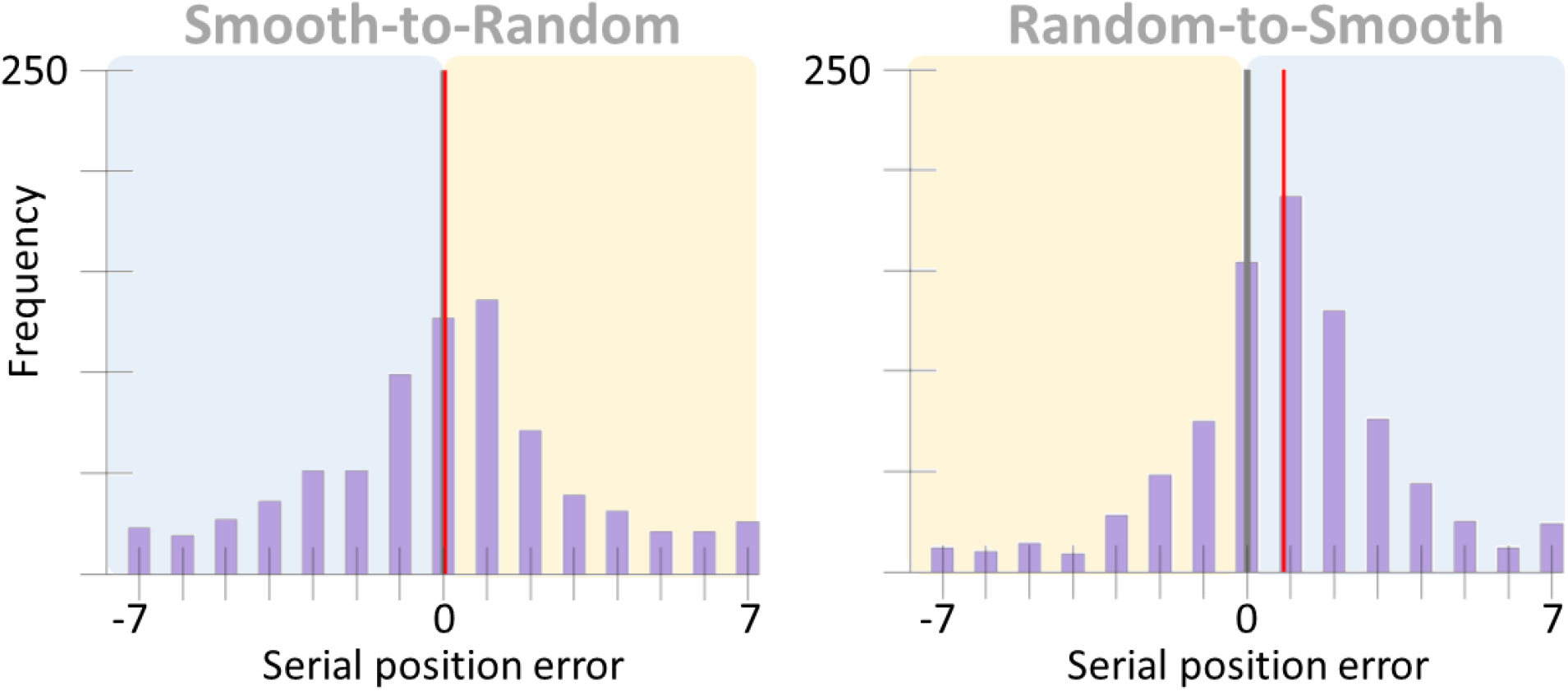
Experiment 3’s results. The x-axis represents serial position error had the color been presented as smoothly changing both before and after the cue. For the half of the graph highlighted in blue, this trajectory matched what the participant was shown. The portion of the graph highlighted in yellow is the error compared to a presentation that *was not shown to participants* but was extrapolated by continuing the smooth trajectory presented. For each condition the grey vertical line marks the position of the cued color. The red vertical line marks the condition mean.

On average, per participant, 11 trials (*SD* = 4) from the Smooth-to-Random condition and 7 trials (*SD* = 4) from the Random-to-Smooth condition were excluded from analysis based on the criteria outlined in Experiment 1. There was a significant delay in selection latency of 88 ms in the Random-to-Smooth condition compared to the Smooth-to-Random condition, *p* < .001, 95% CI [62, 112]. Note that this pattern of results is opposite to that expected from the prediction theory. For completeness, the data is also presented in its raw form (Figure 7). However, as there was no explicit hypothesis for the data in this format, no analysis was performed.

**Figure 7.**
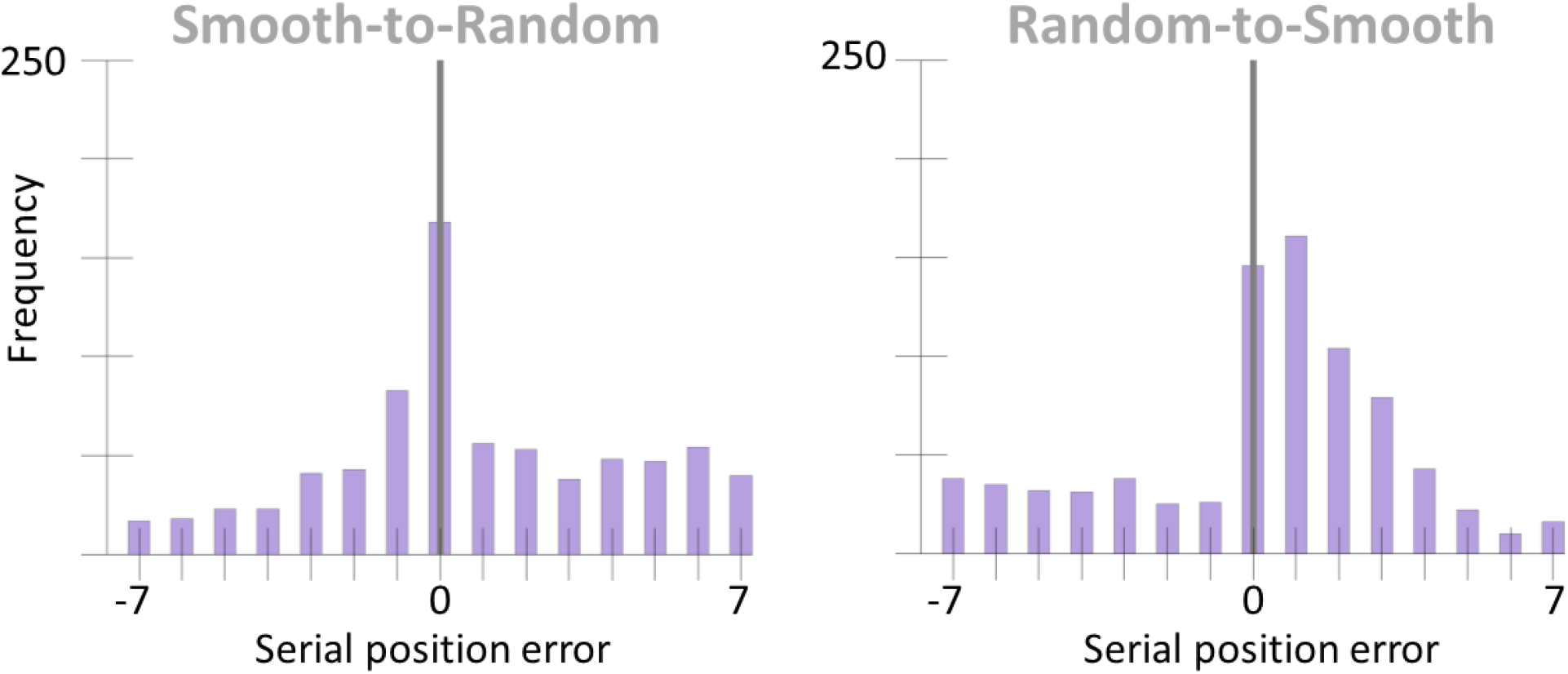
Serial position errors of Experiment 3 comparing reports to actual colors presented to participants (i.e. smooth and random presentation before or after the cue depending on condition). The grey vertical line marks the position of the cued color.

## Experiment 3b

Experiment 3b was run as a replication of Experiment 3.

### Method

An independent sample of 30 undergraduate participants were collected. The same method for recruiting and obtaining informed consent was used in this experiment as described in previous experiments. The exact same stimuli and procedure was used in Experiment 3b as was used in Experiment 3.

### Results

The same analysis methods described in Experiment 3 were used in Experiment 3b. Twelve trials (*SD* = 5) were excluded from analysis for the Smooth-to-Random condition on average while 9 trials (*SD* = 5) were excluded on average from the Random-to-Smooth condition. Again, there was a significant delay in selection latency of 47 ms in the Random-to-Smooth condition compared to the Smooth-to-Random condition, *p* <.001, 95% CI [47, 75] (Figure 8). As in Experiment 3, the data is also presented in its raw form though no analysis was conducted on it (Figure 9).

**Figure 8.**
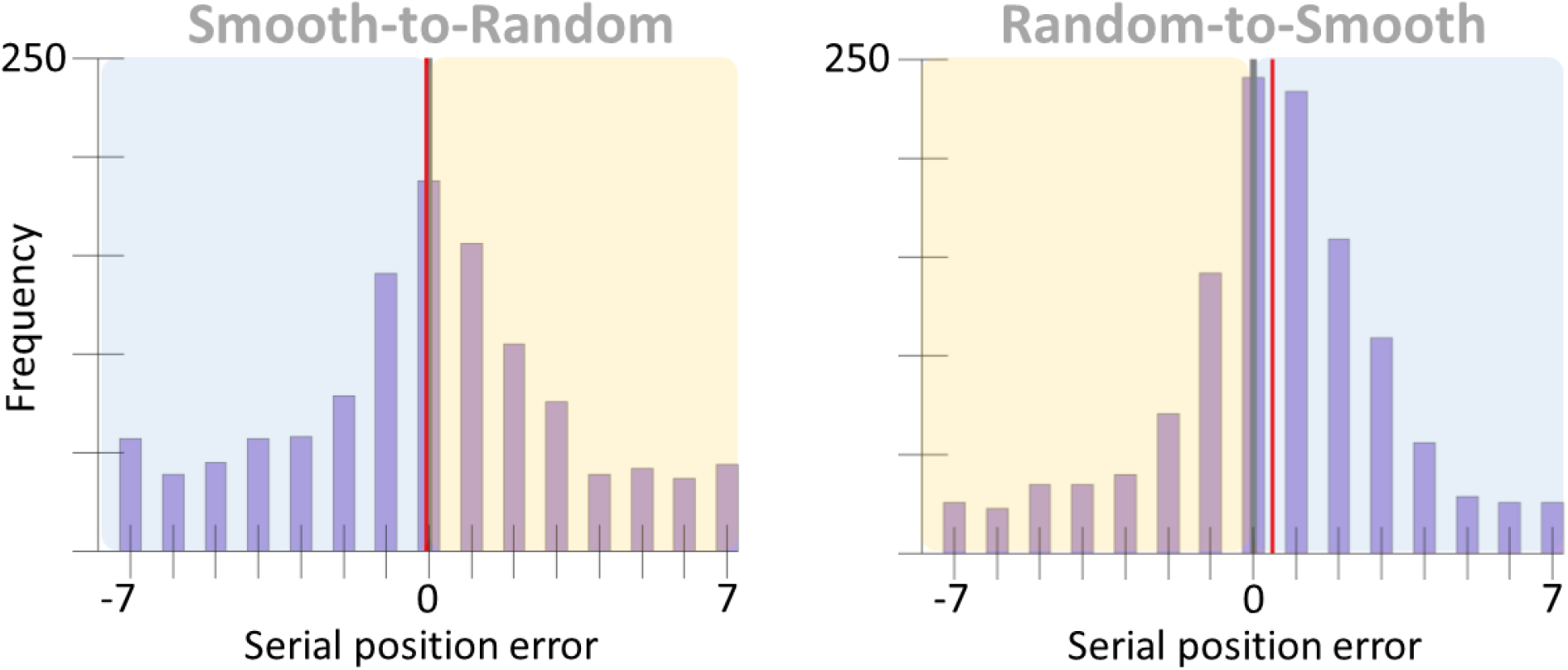
Results from Experiment 3b. As in Experiment 3’s graph, the x-axis represents serial position error had the color been presented as smoothly changing both before and after the cue. The blue box highlights the portion of the trajectory that was shown to the participant. The yellow box highlights the portion of the trajectory that *was not shown to participants* but was extrapolated by continuing the smooth trajectory presented. The grey vertical line marks the position of the cued color and the red vertical line marks the condition mean in both graphs.

**Figure 9.**
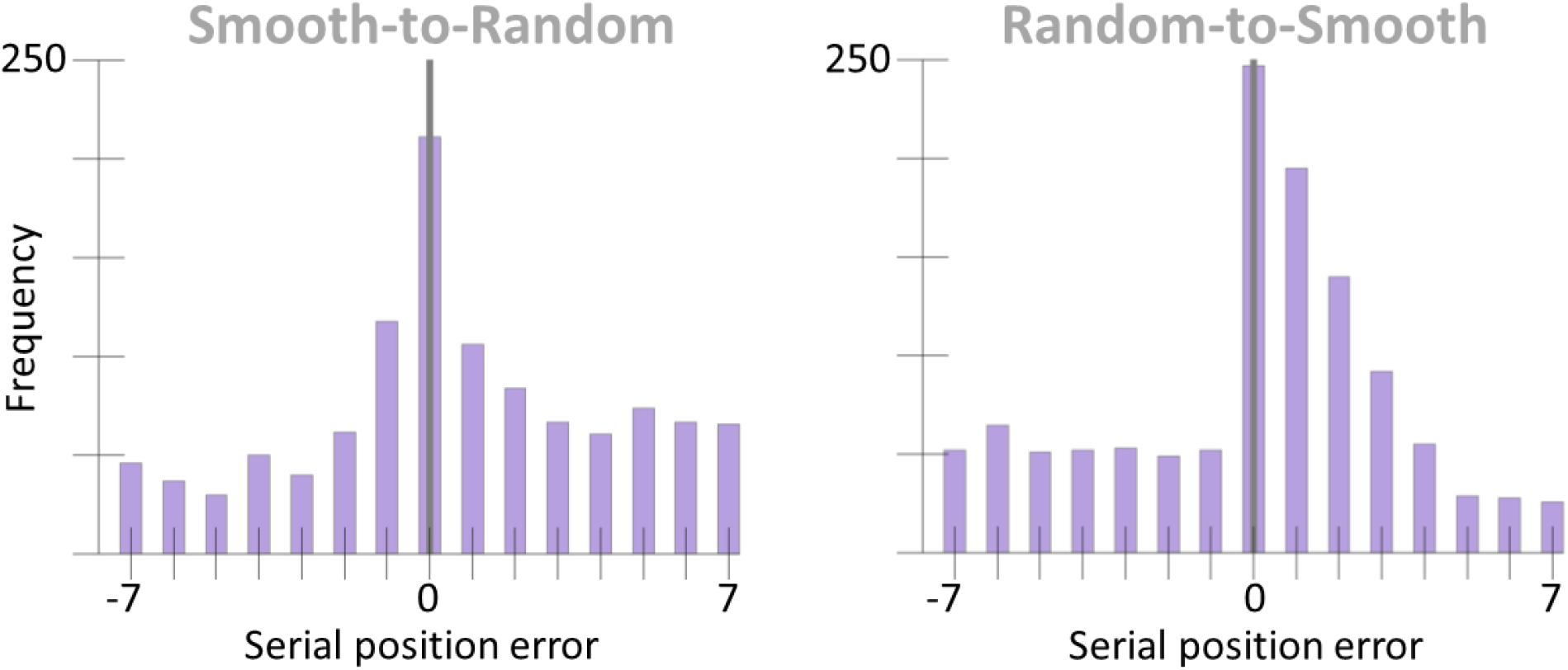
Serial position errors of Experiment 3b comparing reports to actual colors presented to participants (i.e. smooth and random presentation before or after the cue depending on condition). The grey vertical line marks the position of the cued color.

### Discussion of Experiment 3 and 3b

The results from both Experiment 3 and 3b support the idea that the later selection latency seen in the Smooth condition compared to the Random condition in Experiments 1 & 1b is not caused by the smoothness presented prior to the cue (as is assumed by prediction-based theories) but rather by the smoothness presented after the cue. However these results do not rule out the role of prediction entirely but rather one way it might have been used. This will be discussed further in the General Discussion.

## Experiment 4

As a further test of the effect of predictability of the color trajectory, in the present experiment the trajectory was made more similar across trials by presenting the exact same color trajectory every trial. This was done by using only a single color ring, with the same starting color value and direction of change on every trial. The parameters that changed trial-to-trial then was what time the cue appeared and around which disk was it presented. If predictability leads to a later selection latency, this practiced trajectory should produce a later selection than seen in the Smooth_fine condition of Experiments 2 & 2b.

### Method

#### Participants

Seventeen undergraduates, age 18-23 with normal or corrected to normal vision, were used for this experiment with the same recruitment and consent procedure as outlined in Experiments 1, 2, & 3.

#### Stimuli & Apparatus

The same stimuli used in Experiments 1, 2 & 3 were used in this experiment. Color ring 1 was the only color ring used for this experiment (see Methods of Experiment 1 for details)

#### Procedure

A similar procedure was used in Experiment 4 as was used in Experiment 2. Again, participants fixated on a cross in the middle of the screen while monitoring two changing color disks. This time, both disks used the same color ring trajectory. At the start of every trial the disk on the left started its trajectory at the 1° (RGB values: [255, 120.28, 172.2]) color while the disk on the right started at the 180° color (RGB values: [0, 198.1, 168.8]). Both disks moved through their trajectories in the same direction on every trial. The side and time of the cue was randomized as in the previous experiments. As in the Smooth_fine condition of Experiment 2, the colors updated at a rate of 4 degrees every 27 ms. The reporting method used was identical to that in the previous experiments. There was only one condition in this experiment with a total of 88 trials.

### Results

**Figure 10.**
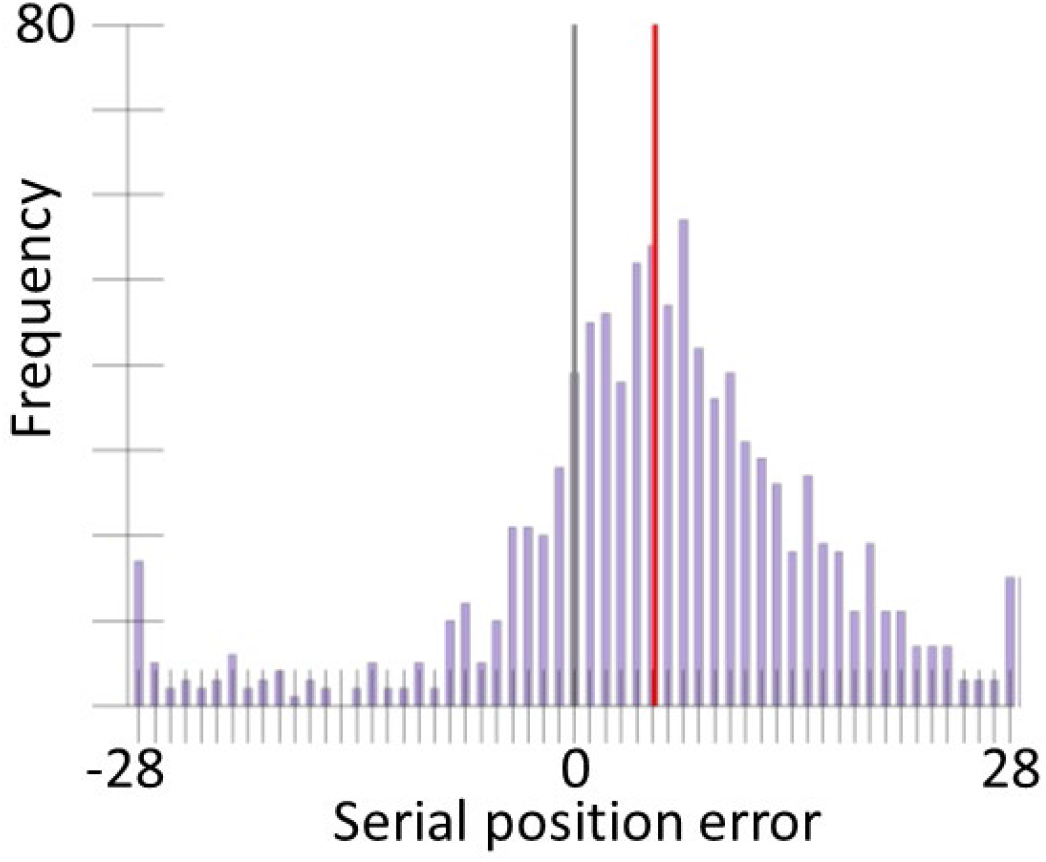
Temporal error distribution from Experiment 4. The grey line indicates the position of the cued color (position zero) and the red line indicates the mean of the distribution.

As in Experiment 2 and 2b the 8 degree criterion was used for excluding trials due to the similarity of colors presented from one time point to the next. This resulted in an average of 29 trials excluded per participant (*SD* = 12). The selection latency of this experiment was 25 ms earlier than that of the Smooth_fine condition in Experiment 2 and 49 ms earlier than that of the Smooth_fine condition in Experiment 2b. As the hypothesis was whether the predictable trajectory would lead to a later selection latency, which was not the case given that, if anything, it is earlier, no analysis was performed on this data.

### Discussion

In Experiments 1, 2, & 3, four different color rings were used and the starting color and direction of change were randomly selected on each trial. This was done to try to avoid potential effects of participants learning the color trajectory. In this experiment, it was much easier to learn the color trajectory and yet the delay was not increased relative to the Smooth_fine condition of Experiments 2 & 2b, reinforcing the conclusion from Experiment 3 that predictability of the sequence is not the cause of the latency to report a color.

## General Discussion

The experiments presented here found that the more smoothly the color of the disk changed, the later the reported color value was. However this was only true if this smooth change occurred after the cue onset. Like most empirical results, these findings have several potential explanations. We highlight our attentional drag theory, in which the smooth change of a feature keeps attention engaged for longer than a condition containing random transitions in feature space. This explanation has the virtue of simplicity, and has also been formally defined with a neurocomputational model. The attentional drag theory provides an explanation for why visual transients are such an effective visual stimulus, not just for capturing attention, but also for disengaging it.

The luminance transients that accompany sudden visual changes have long been suggested to capture or engage attention (Jonides & Yantis, 1988), and sometimes to disengage attention or control binding (Holcombe & Cavanagh, 2008; Holcombe, 2009; Fujisaki & Nishida, 2010). Such findings have typically been explained by proposals that transients directly affect attention – for example, that temporally high-pass cells have a particularly strong connection to neurons that elicit orienting (Motoyoshi & Nishida, 2001). We do not contest such theories here. However, our model does explain how the *disengagement* of attention may be a result not of transients per se but rather because transients are highly associated with the autocorrelation of features that results in lasting featural activity in our model. Future work may be able to tease these concepts apart, perhaps by superposing luminance transients on an otherwise smoothly changing stimulus.

Our core finding was shown by Experiments 1 & 2, documenting a relationship between selection latency and feature change, while Experiment 3 ruled out the possibility that this later selection was the product of future prediction based on a predictable trajectory presented prior to the cue. Experiment 3 showed that in the Smooth-to-Random condition participants did not report future colors along the trajectory when the color changed randomly after the cue. In contrast, when the colors before the cue did not have a consistent trajectory (the Random-to-Smooth condition), participants did have a long selection latency. The results of Experiment 4 provided further evidence against prediction being involved as the sequence was even more predictable but selection latency was not further delayed. A possible alternative explanation for Experiment 3’s results is that color in the smooth presentation are easier to perceive and so reports are biased towards colors presented in the smooth portion of the trial, rather than attention being dragged along in time, as proposed here. However such an explanation would not account for the increased latency in Experiment 1’s Smooth condition where colors are presented smoothly throughout the trial and therefore should produce no bias. The attentional drag hypothesis provides a single explanation for the entire set of observed results.

Functionally, the processing underlying the attentional drag theory may, in effect, capitalize on the statistics of the visual environment. The objects present in our environment typically change slowly or smoothly, so if our principal interest is particular objects, once allocated to a particular object, attention should on average remain there for a while. The sudden change that can disengage attention then helps to temporally segment the scene into distinct objects, which may be complementary to the brain’s parsing of event information on much larger timescales (Zacks, Speer, Swallow, Braver, & Reynolds, 2007; Richmond & Zacks 2017).

### A computational model of the attentional drag

A computational model was developed to explain how the temporal engagement of attention could be extended by presenting a sequence of smooth feature changes.

The model is based on the idea that attention is a recurrent process, as described in a previous model of attentional fluctuations over time (i.e. eSTST, Wyble, Bowman & Nieuwenstein, 2009). The eSTST model simulates attention as part of a positive feedback loop in which incoming sense data (the cue) triggers the deployment of attention at a given spatial location. Attention then amplifies feedforward processing at that location to facilitate the encoding of information into memory. Due to the recurrent nature of attention, this amplification in turn prolongs the deployment of attention, leading (with typical RSVP stimuli) to brief attentional episodes on the order of 150ms in duration, which explains the prevalence of effects like lag-1 sparing in the attentional blink.

The new model proposed here simply elaborates on the input neurons of this architecture by giving them overlapping tuning curves in a given feature domain (e.g. color, orientation). With the addition of such tuning curves, the same model architecture gives rise to the attentional drag phenomenon. A smoothly changing color trajectory causes more activation of each individual color neuron than in the random condition, and that greater activation boosts the activity of an attention neuron over its activation in the random condition, prolonging the engagement of attention (Figure 13). To be clear, this model is not however just the eSTST model with an elaborated stimulus representation. It has the same basic architecture as the eSTST model, but it does not include memory encoding processes, it uses more biologically plausible dynamics, and it runs at a finer time step.

**Figure 11.**
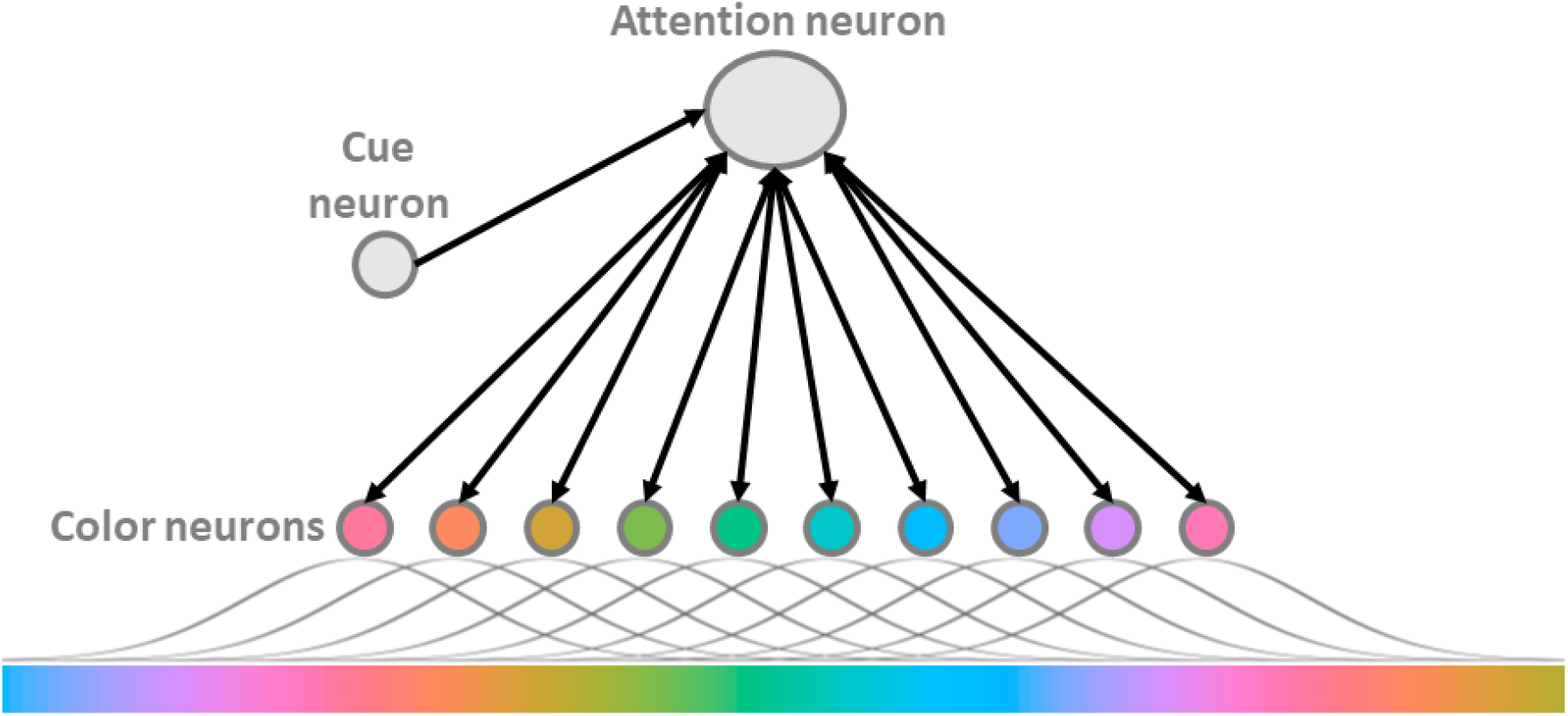
A schematic of the model used to explain the attentional drag theory. Black arrows indicate excitatory connections. For illustration purposes only 10 color neurons are depicted but in the model there are 360. Each neuron is receives input from presented colors under a Gaussian distribution. When the cue neuron is triggered it excites the attention neuron over threshold which triggers the excitatory feedback loop between the attention neuron and the

**Figure 12.**
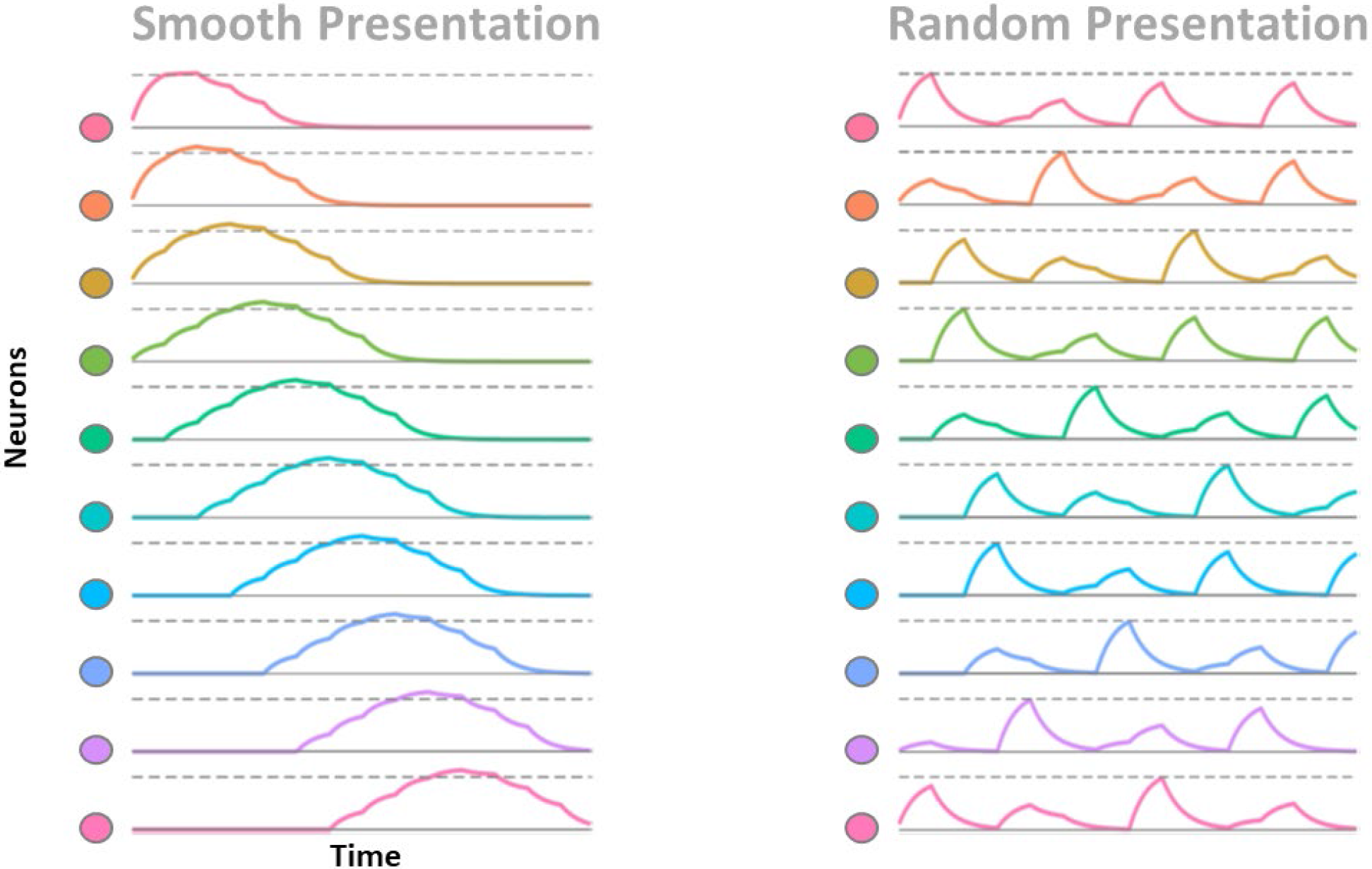
Color neuron activation over time for the Smooth (left) and Random (right) conditions in the absence of attentional amplification. The solid grey line in each graph indicates zero while the dotted grey line indicates the level of activation reached by a color neuron when its optimal color is presented for a 150ms (the SOA used for the Random and Smooth conditions in Experiments 1 & 3). In the Smooth condition, color neuron activation builds on itself as similar colors are presented sequentially. Conversely in the Random condition, no such accumulation of activation is available as the shift in colors presented from one time step to the next is too great. Note that the colors used in the illustration of the neurons and their activation are made to be visually distinctive for clarity and do not represent the color distance between neurons in the actual model.

**Figure 13.**
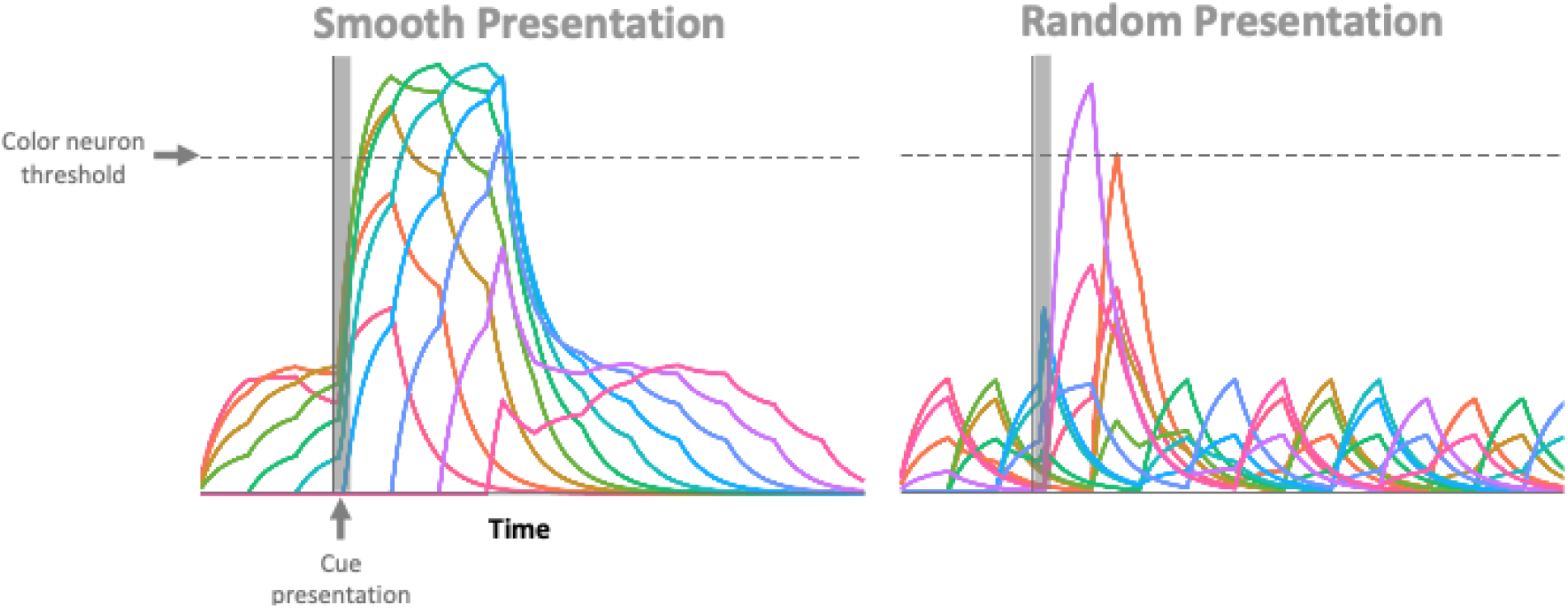
Color neuron activation levels. The grey transparent box denotes the time in which the cue was present on the screen. In the smooth presentation simulation, when attention is triggered in response to the cue, the excitatory feedback loop is extended in time resulting in more color neurons excited over threshold. Conversely in the random presentation simulation much fewer neurons are excited over threshold. Note that the colors used in the illustration of the neurons and their activation are made to be visually distinctive for clarity and do not represent the color distance between neurons in the actual

To understand the drag phenomenon in more detail, consider that each presented color as activating a distributed population of neurons. Smooth changes in color result in the sequential activation of overlapping neural representations of color. This carry-over of activation from one time step to the next interacts with the processes of attentional activation in this model to increase selection latency. Our computational model simulates neuronal membrane potential dynamics with differential equations (O’Reilly & Munakata, 2000), but should be viewed as a simple abstraction – it has only three main components: an array of color sensitive neurons, a cue sensitive neuron, and an attention neuron (Figure 11). Color is represented as a vector representing the activation of 360 stimulus-responsive neurons with Gaussian-shaped tuning curves, which very roughly approximates V1 cells (Wachtler, Sejnowski, & Albright, 2003). In fact V1 cells favor certain directions in color space, whereas we assume uniform tuning across L*a*b space. Future studies could systematically investigate this to assess whether the color representation most resembles the representation in V1 or another neural population. Therefore, any one color evokes a gradient of activation across the population of 360 neurons. A consequence is that when similar colors are presented sequentially (as in the Smooth condition) each color neuron is active for a relatively long period of time, due to the colors presented both before it and the colors presented just after it contributing to its activation (Figure 12). Conversely, when colors are presented in random order, the color presented at one time step is less likely to have overlapping activation with the color presented in the previous and following time steps, resulting in color neurons only receiving excitation for the presentation duration of their preferred color. In summary, smooth feature changes result in higher activation across the population of sensory neurons.

By itself, the higher sensory activation resulting from smooth color changes is not enough to explain the later apparent sampling time observed in the data. This emerges from the effect of the higher sensory activation on the attentional selection mechanism.

It is widely accepted that reporting a stimulus involves feedback from higher-order neurons to sensory neurons (Saalman, Pigarev, & Vidyasagar, 2007), which in our model is accomplished by the “attention neuron”, which is triggered by a cue-responsive neuron. It is the feedback loop between the attention neuron and the sensory neurons that ultimately causes prolonged attentional engagement in the presence of the increased activation caused by overlapping tuning curves with a temporally autocorrelated stimulus.

Specifically, when the cue is presented, this cue neuron is excited over threshold and sends excitation to the attention neuron. Once the attention neuron has been excited over threshold, it amplifies the excitation being received by all of the color neurons. For color neurons which are currently not receiving excitation (i.e. the color they are responsive to is not being presented) this attentional amplification does nothing. However for colors that are responsive to the current input, this attentional amplification excites them over threshold, allowing them to excite the attention neuron in turn. Thus, the presentation of the cue triggers the deployment of attention and consequently puts the system into an excitatory feedback loop. In the case of randomly presented colors, this loop has a short duration, decaying quickly after the offset of the cue. However when the system is presented with a smoothly changing color stimulus, there is overlap in the activation of color neurons from one time step to the next, which causes increased excitation to the attention neuron. Therefore, when the cue neuron triggers the deployment of attention, this feedback loop is sustained for longer in the Smooth condition compared to the Random one (Figure 13). The longer the duration of this excitatory loop between the attention and color neurons, the more color neurons that are excited over threshold (Figure 14). An additional experiment, included in the Supplement, supports the model by showing this phenomenon is robust to different cue durations (Experiment S1).

**Figure 13.**
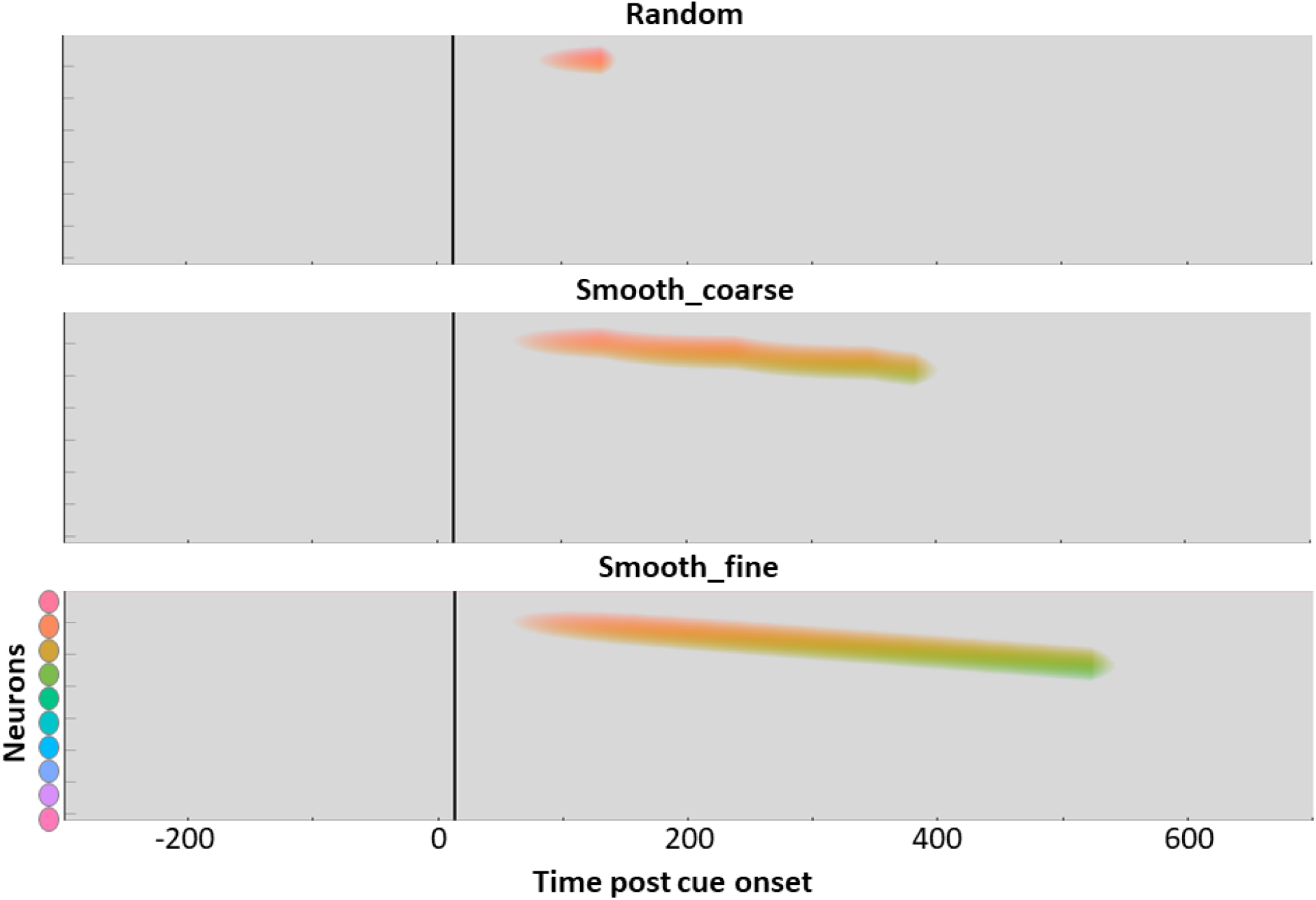
Illustration of the attentional drag effect on color neuron activation in the Random, Smooth_coarse and Smooth_fine conditions. The y-axis is the 360 color neurons and the x-axis is the time post cue onset. The black line in in each graph marks the time at which the attention neuron crossed threshold. The non-grey pixels of the graph mark color neurons over threshold in time.

While this model does not make specific claims as to how the ultimate color is selected for report, the extended activation means greater activation for later-occurring colors and leads to a higher probability of reporting a later color in the Smooth condition compared to the Random condition. One possibility is that the ultimate selection from the extended color representation in the Smooth condition is made by an endogenous form of attention that has been shown to have slower response times (Müller & Rabbitt, 1989) and is poor at selecting information with a short latency from a dynamic stimulus (Holcombe & Cavanagh 2008).

The model simulation matches the pattern of results seen in Experiments 1 & 1b – the Smooth condition shows a longer selection latency than the Random condition. The increased color similarity of successive time steps of Experiments 2 & 2b, by boosting the amount of overlap in activated neural representations in the model resulted in later selection latency for the Smooth_fine condition compared to the Smooth_coarse. Without specifying a method of color selection, the model provides a formalized account of the attentional drag theory here and simulates the pattern of results seen in the empirical data. It is important to note that the model does not make any distinction between whether smoothness extends attentional engagement at a location or if a transient in color disengages attention, as these are two sides of the same coin in terms of the neural activation in the model. Finally, a basic property of the model is that the increase in selection latency results from the degree of smoothness presented after the cue. Experiment 3 provided evidence that this is indeed the critical variable, thus ruling out an alternative account based on sampling from predictions of an internal model.

This is an abductive model in that it accounts for the empirical results with a simple set of neural mechanisms that are plausible inasmuch as they extend a model that has had success with other attentional phenomena (Wyble, Bowman & Nieuwenstein, 2009). However, previous research has had a different interpretation of similar findings. Namely, Sheth et al., (2000) similarly found a long selection latency when reporting the color of a disk at the time of a cue as the disks smoothly changes between red and green. Sheth and colleagues proposed that, as the color changes smoothly, priming occurs, creating a ramp-up in activation of subsequent colors along the trajectory. When attention is triggered by the cue, it resets this build-up of activation and so the selection of color is delayed until priming can once again ramp-up a colors activation sufficiently to be selected. This accounts for their results as well as the selection latency of the Smooth condition presented here. However, this theory, as is, would not explain the pattern of selection seen in the Random condition. Explaining this additional finding requires the priming theory to add some sort of immediate enhancement performed by attention in order to privilege the cued color for selection. Without this additional form of selection, every color presented in the Random condition would have an equal chance of being selected. This attentional enhancement could possibly be incorporated into the theory, in addition to the resetting of priming by attentional shifts, however, as this requires two distinct roles for attention, it could be argued that the attentional drag theory offers a simpler account for the experimental results. Further elaboration of an attentional priming account, ideally with a computational instantiation would be a key step in testing the distinction between these accounts since it is difficult to compare models with several different components without a concrete specification.

### Conclusion

The empirical results presented here support an attentional drag theory wherein gradual or smooth changes at a location maintain attentional engagement compared to a salient feature change which disengages attention. These results demonstrate how feature dynamics affect sensory information extraction. When features make jumps in feature space, attention is able to select information at the time of the cue. This is consistent with previous findings that have shown a transient signal at a location aids in temporal sampling (Holcombe & Cavanagh, 2008; Nishida & Johnston, 2002). Conversely, when features change smoothly in time, selection is delayed. These results suggest that the visual system samples information differently depending on the nature of change of the input. As features often change smoothly in our visual world, understanding this phenomenon could provide important insights into how attention tracks stimuli. For example, keeping attention anchored to stimuli that exhibit smooth changes may provide an inherent form of event segmentation that combines visual information from a sequence of time points only when they are likely to comprise the same visual event or perception of a single object.

## Supplemental

**Table S1.** Analysis for Experiments 1 & 1b, varying threshold. Statistics for Experiments 1 & 1b for 3 different levels of constraints. In “No constraint”, all trial were excluded except those that were tied for minimum distance between two colors presented colors. The “< 16°” version is what is presented in the paper and the “< 6°” version is where all trials outside of 6 degrees of the presented colors were excluded.

|  |  | Mean Difference<br>(ms) | *p*-value | 95% CI of mean difference<br>(ms) |
| --- | --- | --- | --- | --- |
| Exp 1 | < 6° | 91 | <.001 | [54, 129] |
|  | < 16° | 110 | <.001 | [76, 135] |
|  | No constraints | 114 | <.001 | [81, 144] |
| Exp 1b | < 6° | 104 | <.001 | [66, 142] |
|  | < 16° | 113 | <.001 | [78, 137] |
|  | No constraints | 111 | <.001 | [80, 142] |

**Figure S1.**
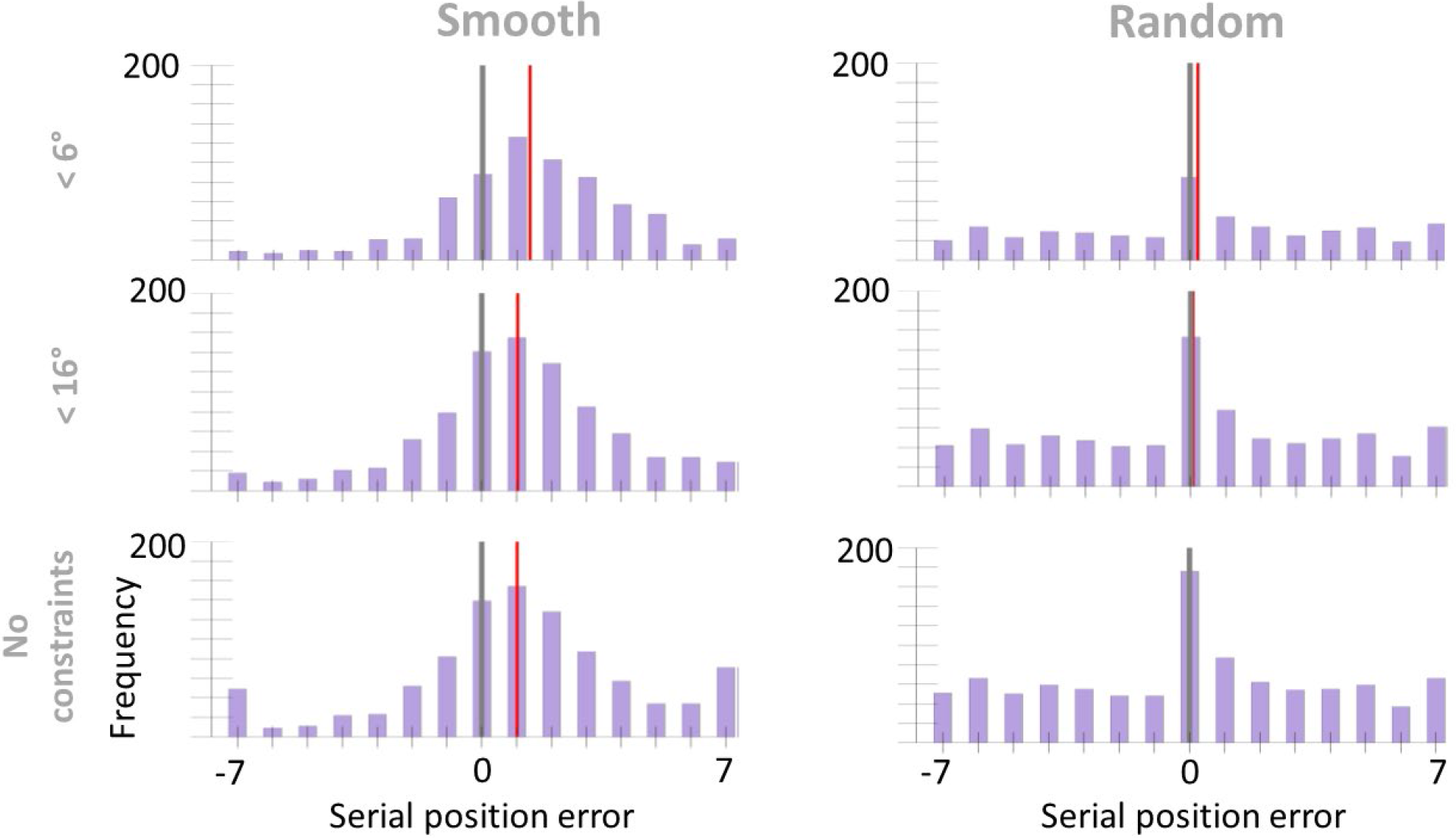
Serial position error histograms for Experiment 1. Top: data analyzed using a threshold of 6 degrees, such that trials with a minimum difference exceeding 6 degrees from the colors presented 7 positions before and after the cue were excluded. Middle: data analyzed using a threshold of 16 degrees (presented in paper). Bottom: data analyzed without constraints, only excluding those trials that had a tied minimum difference between 2 presented colors. The grey vertical line marks the position of the cued color and the red line marks the condition mean.

**Figure S2.**
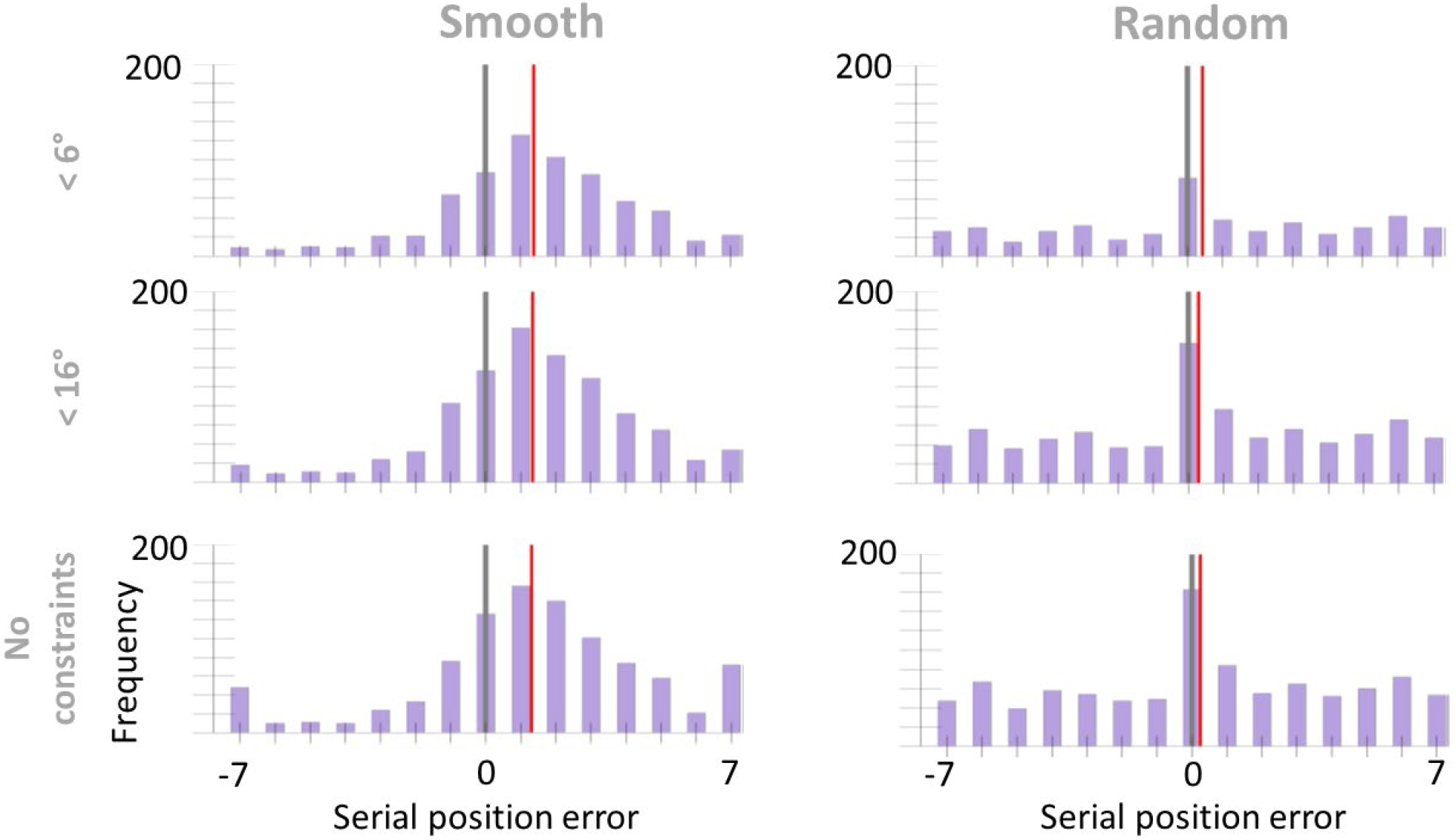
Serial position error histograms for Experiment 1b. Top: data analyzed using a threshold of 6 degrees, such that trials with a minimum difference exceeding 6 degrees from the colors presented 7 positions before and after the cue were excluded. Middle: data analyzed using a threshold of 16 degrees (presented in paper). Bottom: data analyzed without constraints, only excluding those trials that had a tied minimum difference between 2 presented colors. The grey vertical line marks the position of the cued color and the red line marks the condition mean.

**Figure S3.**
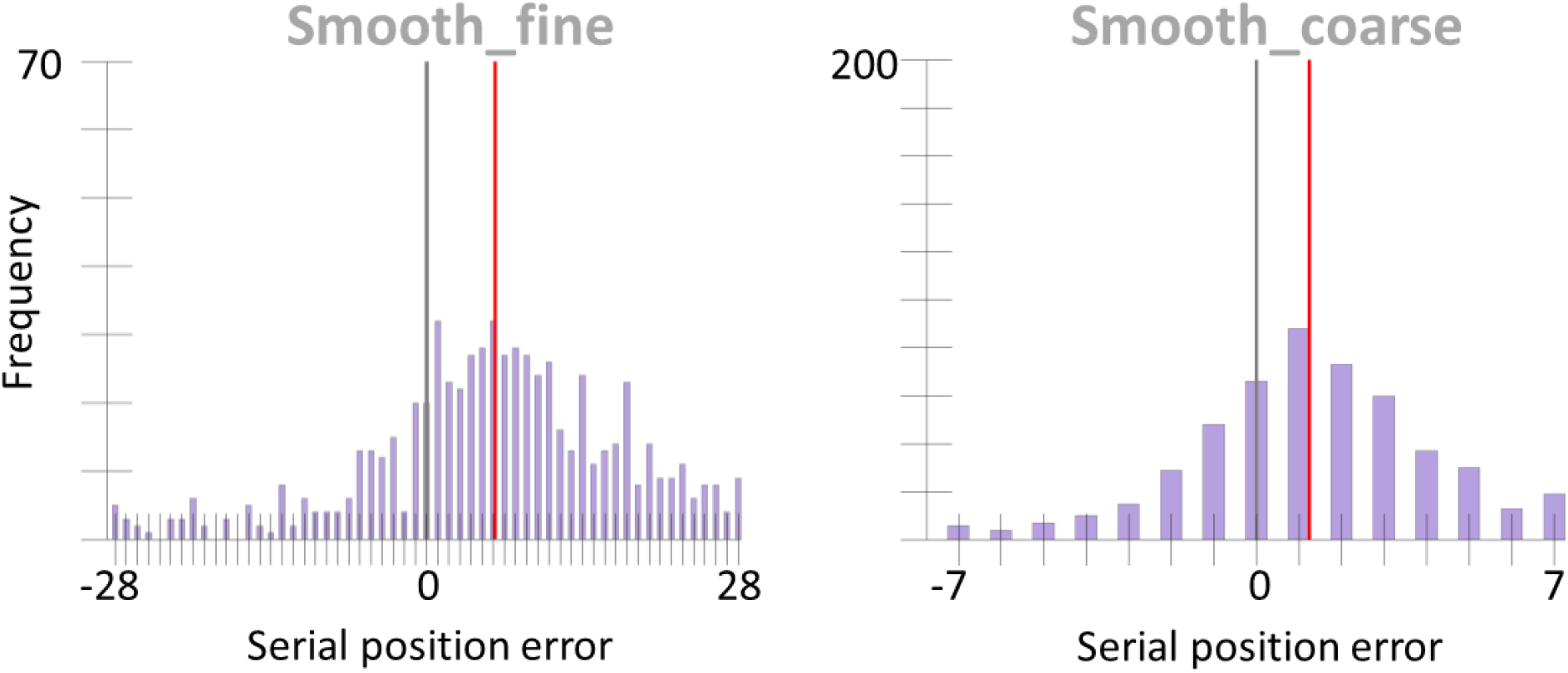
Experiment 2’s data re-analyzed with a lower exclusion criteria. Here, all trials where the reported color value was over 4° from all presented colors were excluded from analysis. This resulted in average of 18 trials (*SD* = 4) excluded from the Smooth_coarse condition and 12 trials (*SD* = 4) excluded from the Smooth_fine condition. There was a significant delay in selection latency of 31 ms in the Smooth_fine condition compared to the Smooth_coarse condition, *p* = .04, 95% CI [4, 59]. The grey vertical line marks the position of the cued color and the red line marks the condition mean.

**Figure S4.**
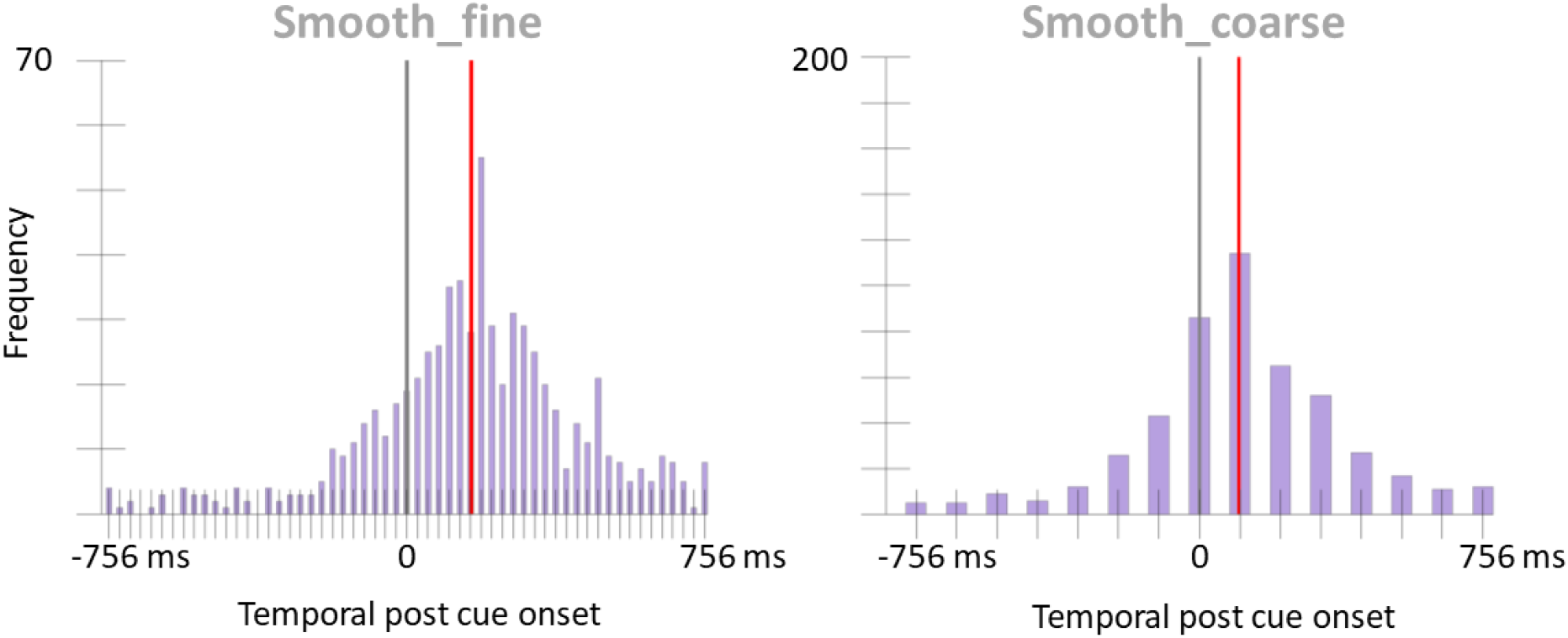
Experiment 2b’s data, excluding all trials where the reported color value was over 4° from all presented colors. Nineteen trials (*SD* = 4) were excluded from the Smooth_coarse condition and 12 trials (*SD* = 3) were excluded from the Smooth_fine condition. There was a significant delay in selection latency of 59 ms in the Smooth_fine condition compared to the Smooth_coarse condition, *p* = .001, 95% CI [32, 84]. The grey vertical line marks the position of the cued color and the red line marks the condition mean.

## Experiment S1

Experiment S1 was conducted to test whether the duration of the cue would have an effect on the selection latency of a feature. In previous unpublished pilot experiments associated with the experiments described in Holcombe & Cavanagh (2008), attentional selection from temporally autocorrelated stimuli seemed more effective with a longer cue. As a brief cue is not well matched to the poor temporal resolution of attention (Holcombe, 2009), with a brief cue it is possible that sampling must occur via more endogenous attention, resulting in longer latency. This suggests that with a longer cue, there might be a shorter sampling latency.

The attentional drag theory, in contrast, attributes long selection latencies to the temporal autocorrelation of smoothly changing stimuli and does not suggest there should be an effect of duration of the cue. To test this, two conditions were compared, varying the duration of cue presentation.

### Method

#### Participants

A sample of 24 undergraduates age 18-23 with normal to correct to normal vision were used for this experiment with the same recruitment and consent procedure as outlined in all of the previous experiments.

#### Stimuli & Apparatus

The same stimuli used in the first 3 experiments were used in this experiment.

#### Procedure

In this experiment participants again maintained fixation on a cross in the middle of the screen and monitored two changing colored disks. The colors changed in the same fashion as the Smooth condition of Experiment 1, stepping 16 degrees around a color ring every 108 ms. Participants were told to report the color of the disk at the time of the cue. The same cueing and reporting method used in the previous 3 experiments were used here.

In this experiment there were two conditions. The Short condition was identical to the Smooth condition of Experiment 1 where the cue was presented only for the first 27 ms of the cued color’s presentation. In the Long condition the cue was presented for 108 ms (the same duration as the cued color).

### Results

The same permutation analysis and bootstrap method described in Experiment 1 were used here. There was a non-significant delay in the selection latency for the Long Cue condition compared to the Short Cue condition (24 ms), *p* = .08, 95% CI [-3, 50]. If anything, the data suggest that the latency is longer in the Long condition, which is contrary to our concern that the Long condition might conceivably reduce or eliminate the attentional drag effect, if the effect reflected ineffectiveness of attention specific to an overly brief cue.

**Figure S3.**
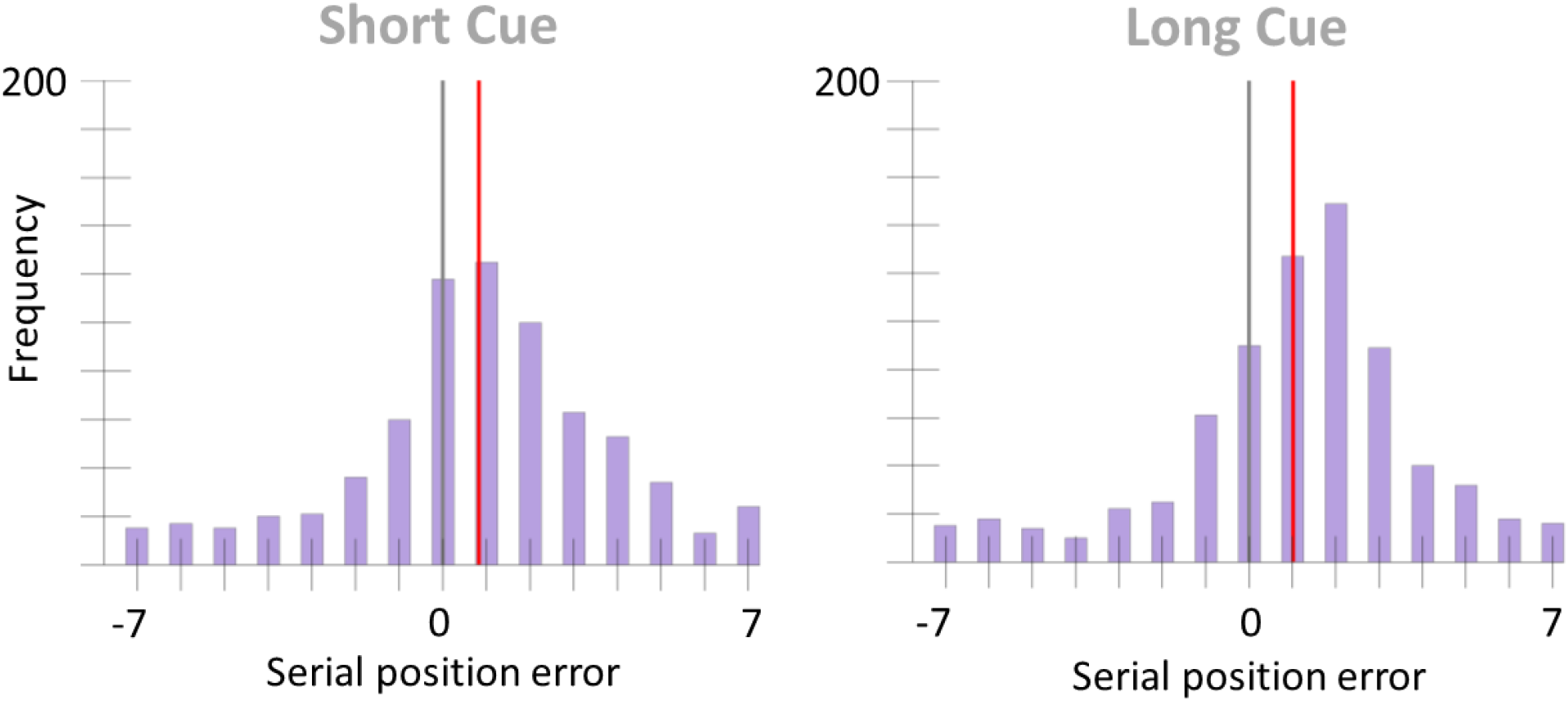
Serial position error histogram for Supplemental Experiment 1. The grey vertical line marks the position of the cued color and the red line marks the condition mean.

## Footnotes

1 This analysis was also repeated with permutations at the subject level. Here the mean difference between conditions for each subject was calculated. There was a .5 chance for each subject that this difference was calculated by subtracting the mean of Smooth trials from the mean of Random trials or vice versa. The average of across all of these subject differences was then calculated. This process was repeated 10,000 time to build a distribution of average condition differences, generated under the assumption that there is no true difference between conditions (the null distribution). The true grand average was then compared to this distribution in order to calculate a p-value. The dichotomous outcomes of all tests throughout the paper were identical when using this method and the p-values were of similar magnitude.

## Notes

#### Summary of Updates

edited reference list

https://osf.io/hujwb/

